# Chromatin architecture mapping by multiplex proximity tagging

**DOI:** 10.1101/2024.11.12.623258

**Authors:** Axel Delamarre, Benton Bailey, Jennifer Yavid, Richard Koche, Neeman Mohibullah, Iestyn Whitehouse

**Affiliations:** Memorial Sloan Kettering Cancer Center, Molecular Biology Program; New York, USA; Memorial Sloan Kettering Cancer Center, Center for Epigenetics Research; New York, USA; Memorial Sloan Kettering Cancer Center, Integrated Genomics Operation; New York, USA

## Abstract

Chromatin plays a pivotal role in genome expression, maintenance, and replication. To better understand chromatin organization, we developed a novel proximity-tagging method which assigns unique DNA barcodes to molecules that associate in 3D space. Using this method – Proximity Copy Paste (PCP) – we mapped the connectivity of individual nucleosomes in Saccharomyces cerevisiae. By analyzing nucleosome positions and spacing on single molecule fibers, we show that chromatin is predominantly organized into regularly spaced nucleosome arrays that can be positioned or delocalized. Basic features of nucleosome arrays are generally explained by gene size and transcription. PCP can also map long-range, multi-way interactions and we provide the first direct evidence supporting a model that metaphase chromosomes are compacted by cohesin loop clustering. Analyzing single-molecule nuclease footprinting data we define distinct chromatin states within a mixed population to show that non-canonical nucleosomes, notably Overlapping-Di-Nucleosomes (OLDN) are a stable feature of chromatin. PCP is a versatile method allowing the detection of the connectivity of individual molecules locally and over large distance to be mapped at high-resolution in a single experiment.

## Main Text

Understanding of chromatin structure and how nucleosomes are organized on DNA has advanced rapidly with the utilization of genomics methods that map the products of chromatin digestion by nucleases (1). The high degree of organization and consistent positioning of nucleosomes near important regulatory DNA elements – such as gene promoters – is a pervasive feature of eukaryotic genomes (2–4). Beyond the primary organization of nucleosomes, the folding and long-range interactions of chromatin within the nucleus has profound impact on the regulation of basic DNA transactions, such as gene transcription, DNA replication and repair (5–7). Mapping how chromatin folds in 3D space is typically accomplished with 3C based methods which use pairwise DNA ligation to preserve long-range interactions (8–11). Despite these advances, we have a limited understanding of how nucleosomes are organized across gene bodies, or how multiple regions of chromatin may cluster in 3D space.

Gaps in understanding underscore the limitations of genomics methods that map the average properties of molecules from mixed populations. Indeed, most epigenomics methods capture a tiny fraction of the available information in a sample and cannot simultaneously assay multiple factors. These deficiencies have spurred efforts to develop approaches that map the co-association of DNA, RNA or proteins in crosslinked chromatin with single-molecule precision (12, 13). Methods such as SPRITE and ChIA-Drop offer a significant advance in mapping higher-order interactions in genomes, but are complex and require a crosslinked genome to be broken into smaller pieces by sonication before they are indexed. Unfortunately, sonication fundamentally limits the assay as connectivity between molecules is progressively lost as shearing – hence resolution – is increased. Thus, high resolution assays - such as those involving nucleosome-nucleosome interactions are unlikely to be achieved with methods that separate and index fragmented chromosomes.

Proximity ligation strategies such as Micro-C and Hi-CO (9, 11, 14, 15) can map spatial relationships between nucleosomes without the need for sonication and have revealed important details of chromatin architecture *in vivo*. Yet the reliance of pairwise DNA ligation to preserve 3D contact information heavily biases the readout towards very short-range contacts and multi-way interactions cannot be accessed without use of long-read sequencing (16). Alternative methods using DNA methylation foot-printing with single molecule sequencing can map locations of arrayed nucleosomes on individual chromatin fibers (17–19). Yet such approaches lack the throughput and resolution offered by short-read sequencing of nuclease digestion products, and 3D connectivity of individual nucleosomes cannot be deduced.

Beyond epigenomics methods, contacts between proteins or nucleic acids can also be detected in fixed cells with templated ligation of DNA circles to facilitate detection or sequencing of an amplified circle(20, 21). Alternatively, contacts between a known “bait” protein and its interactors can be mapped by proximity labelling methods, that typically involve tethering enzymatic domains to the bait protein. The tethered enzymes convert small molecules into reactive, diffusible tags which covalently modify molecules near the bait. Such approaches have found broad utility in mapping interactions between known proteins and their contacts. However, the use of an invariant reactive molecule (e.g. biotin (22)) as a diffusible tag limits the utility of these methods to study the interactions of a single bait in each experiment. Thus, multiplex interactions between a diverse array of bait and targets cannot be mapped with conventional proximity labelling methods.

Here, we describe the development of a new technology that employs a novel proximity tagging approach utilizing a diverse array of nucleic acid tags. The use of nucleic acid tags overcomes the limitation of conventional proximity tagging approaches allowing us to uniquely map multiplex interactions between a diversity of baits and targets. Here, we demonstrate the utility of this technology, which we term PCP (Proximity Copy Paste), to investigate chromatin structure from the level of nucleosome organization up to higher-order chromosome folding, using budding yeast as a model. Importantly, by defining the connectivity of individual nucleosomes we show that fuzzy nucleosomes observed in gene bodies in ensemble studies are actually part of larger ordered nucleosome arrays. In addition, by mapping multiplex 3D contacts, we provide evidence that cohesin forms multi-way hubs along chromosomes in G2. Finally, we show that the chromatin remodeling enzyme RSC is responsible for the generation of an overlapping di-nucleosome between divergent gene promoter pairs, demonstrating that alternate nucleosome structures are a stable, intrinsic feature of chromatin *in vivo*. Collectively, we showcase that PCP offers a powerful new approach to map the spatial relationships between molecules.

### A reaction to uniquely tag proximal molecules

Single molecule approaches SPRITE and ChIA-Drop can map spatial relationships between molecules by uniquely indexing fragments of crosslinked chromosomes (12, 13). Yet, the need to fragment and physically separate molecules to be uniquely indexed is cumbersome and significantly limits the ability to perform high-resolution, multiplex analysis. We reasoned that the development of a proximity tagging approach, wherein a vast array of locally produced tagging molecules distinctly label spatially related regions of chromatin, would allow the analysis of chromosome folding while preserving the global structure of chromatin.

We developed a system that produces a library of nucleic acid tags which are engineered to encode a diverse array sequences. The system has two components: “seeds”, which are double stranded DNA fragments comprising a T7 RNA polymerase promoter, a unique molecular identifier (UMI) and an annealing sequence; and “receptors” which are double-stranded DNA molecules with a 5’ single-strand overhang complementary to the annealing sequence in the seed (Fig 1a). Upon the simultaneous addition of T7 RNA polymerase and Reverse Transcriptase, the seed will be transcribed and produce multiple copies of RNA molecules (RNA tags) carrying the UMI matching the seed from which it was produced. The RNA tag will diffuse from the seed and anneal to a complementary receptor sequence where the RNA tag will be used as a template by Reverse Transcriptase. The tagging reaction can be quantitated by DNA sequencing – allowing receptors tagged by each seed to be unambiguously identified.

**Figure 1:**
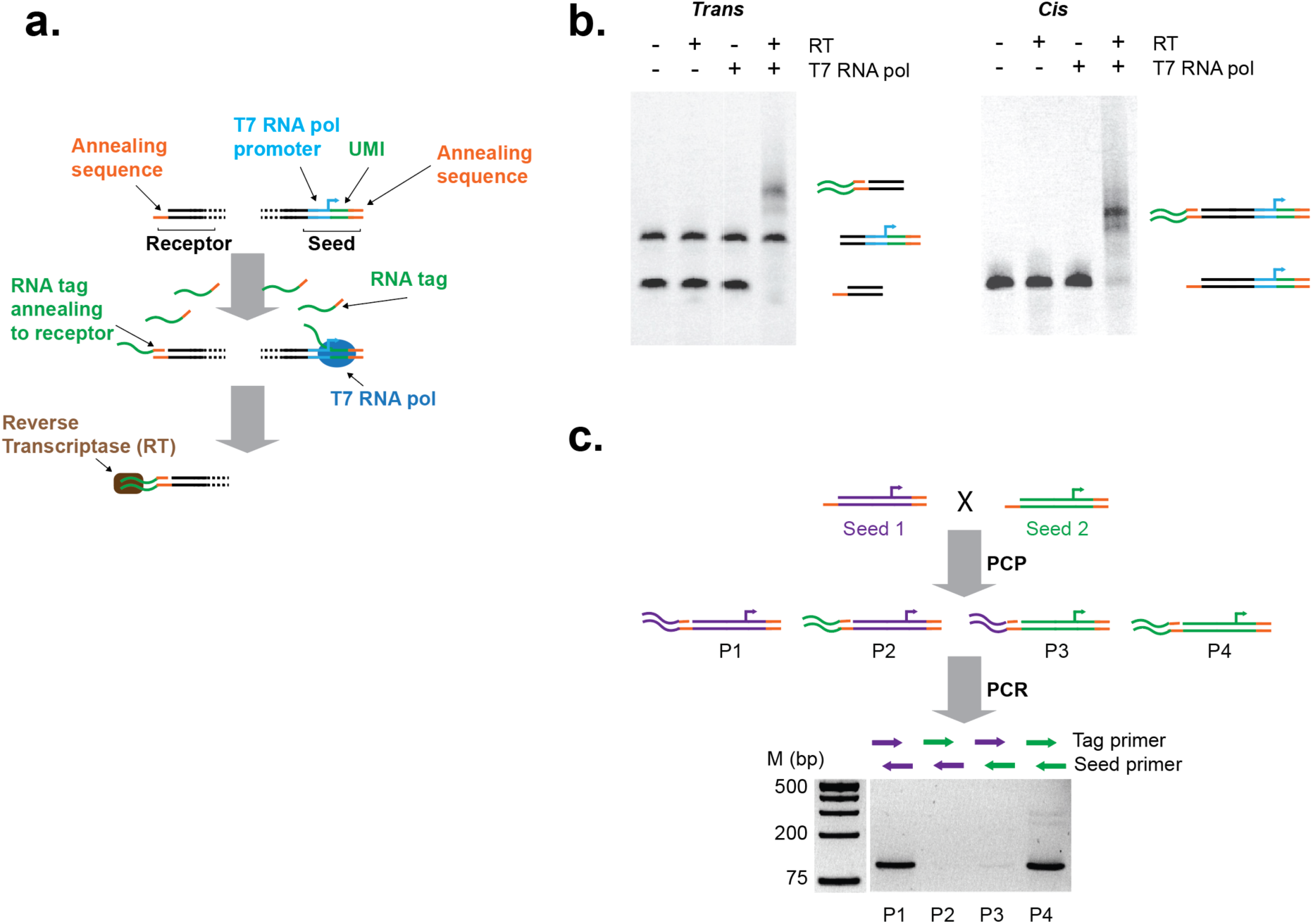
Testing PCP reaction *in vitro.* (**a**) Schematic of the PCP reaction. A seed molecule comprising a T7 RNA polymerase promoter is transcribed to produce an “RNA tag” containing a unique molecular identifier (UMI) and an annealing sequence. The RNA tag diffuses from the seed but can anneal to a complementary annealing sequence on a receptor molecule. The annealed RNA can be used as a template by a reverse transcriptase which will copy the UMI by extending the 3’ end of the receptor DNA molecule. Multiple RNA tags are produced from each seed, allowing multiple receptor molecules to be tagged. (**b**) Migration of PCP products on a native polyacrylamide gel. Left panel, tagging occurs in *trans* when separate seed and receptor molecules are added to the PCP reaction. Right panel, tagging occurs in *cis* if the receptor and the seed are positioned on the two extremities of the same molecule. PCP reactions have high efficiency and occur only in the presence of both polymerases. (**c**) Isolation test of the PCP reaction. Two seeds with receptors in *cis* are mixed in equal proportions. The receptor sequences are identical – 4 products of the PCP reaction are expected if equal mixing and tagging occur (P1-P4). After the reaction, the products can be distinguished by PCR with specific primers (purple or or green arrows). This experiment produces almost exclusively pure products (P1 or P4), highlighting a preference for tagging in *cis*.

Testing the reaction with model templates with simple annealing sequences shows that the PCP reaction works efficiently in *cis* or *trans* with specific elongation of the receptor sequence mediated by the tagging RNA produced from a seed (Fig 1b). Importantly, each seed can produce multiple RNA tags, allowing numerous receptors to be tagged by the same seed. Next, we investigated whether tagging in *cis* or *trans* is favored; two seeds were generated with identical receptors (Fig 1c). The two molecules differ slightly in sequence to allow the products of tagging to be determined by PCR. The two seeds were mixed in equal proportions and incubated in the PCP reaction. Four distinct products can be generated, reflecting whether the tagging reaction occurred in *cis* or *trans* and the fact that a receptor can only be tagged once. Importantly, tagging of receptors in *cis* is markedly favored over *trans* (Fig 1c) – a result that demonstrates that receptors near the seed will be preferentially tagged due to high local concentration of the RNA tag produced from a seed.

### Mapping global chromatin organization

The 3C family of methods uses proximity ligation of DNA to map how chromatin is organized in 3D space (8–10). However, ligation requires direct contact between DNA extremities, whereas PCP could allow the detection of molecules that are proximal but not necessarily ligatable. Moreover, high resolution assays, such as Micro-C can only map pairwise interactions, a limit that PCP can overcome provided that receptors are in excess of seeds. Using *Saccharomyces cerevisiae,* we devised a protocol to use PCP to map 3D interactions in chromatin. For this, we extensively crosslinked chromatin with formaldehyde and DSG before nuclease digestion of the chromatin to the nucleosome level. The crosslinked chromatin is then immobilized on magnetic beads and a mixture of receptors and seeds are ligated to the ends of the nucleosomal DNA. The PCP reaction is subsequently used to tag receptors in proximity to seeds (Fig 2a, Fig S1a). If seeds preferentially tag nearby receptors, PCP should be able to generate a map of how individual molecules (e.g. nucleosomes) associate in 3D space, yet nucleosome level resolution can only be achieved if most of the molecules in the PCP assay can be tagged and detected. Importantly, unlike ensemble epigenomics assays like Micro-C, reaction inefficiencies in PCP cannot be compensated by an increase in input. Through extensive optimization, we found that Micrococcal Nuclease is ill-suited for controlled chromatin digestions as it readily over-digests nucleosomal DNA, rendering the DNA ends poor substrates for end repair and ligation. We reasoned that Caspase Activated DNAse (CAD), which cleaves chromatin in a similar manner to MNase (23), but does not digest nucleosomal DNA (24, 25) and typically leaves extra-nucleosomal DNA after digestion (23), would allow efficient end repair and quantitative ligation of seed and acceptor molecules to crosslinked nucleosomes. Following optimization, we were able to achieve near complete ligation of seeds and receptors to crosslinked chromatin (Fig S1b). The fixed and digested chromatin is ligated with a defined ratio of seeds and receptor molecules (1:10 in the rest of this study) (Fig S1a). Following the 60-minute PCP reaction, chromatin crosslinks are reversed and a library is generated by direct PCR amplification. Sequencing is performed by pair end, allowing insert (footprint) sizes to be accurately mapped (Fig S1c). Each uniquely tagged molecule can be identified by the UMI and the genomic map position of the tagged DNA. We optimized a protocol that utilized ∼100pg of DNA (equivalent to ∼6,000 budding yeast cells) per PCP reaction, allowing us to capture ∼30-50% of the library diversity with moderate number of sequencing reads (Fig S1d).

**Figure 2:**
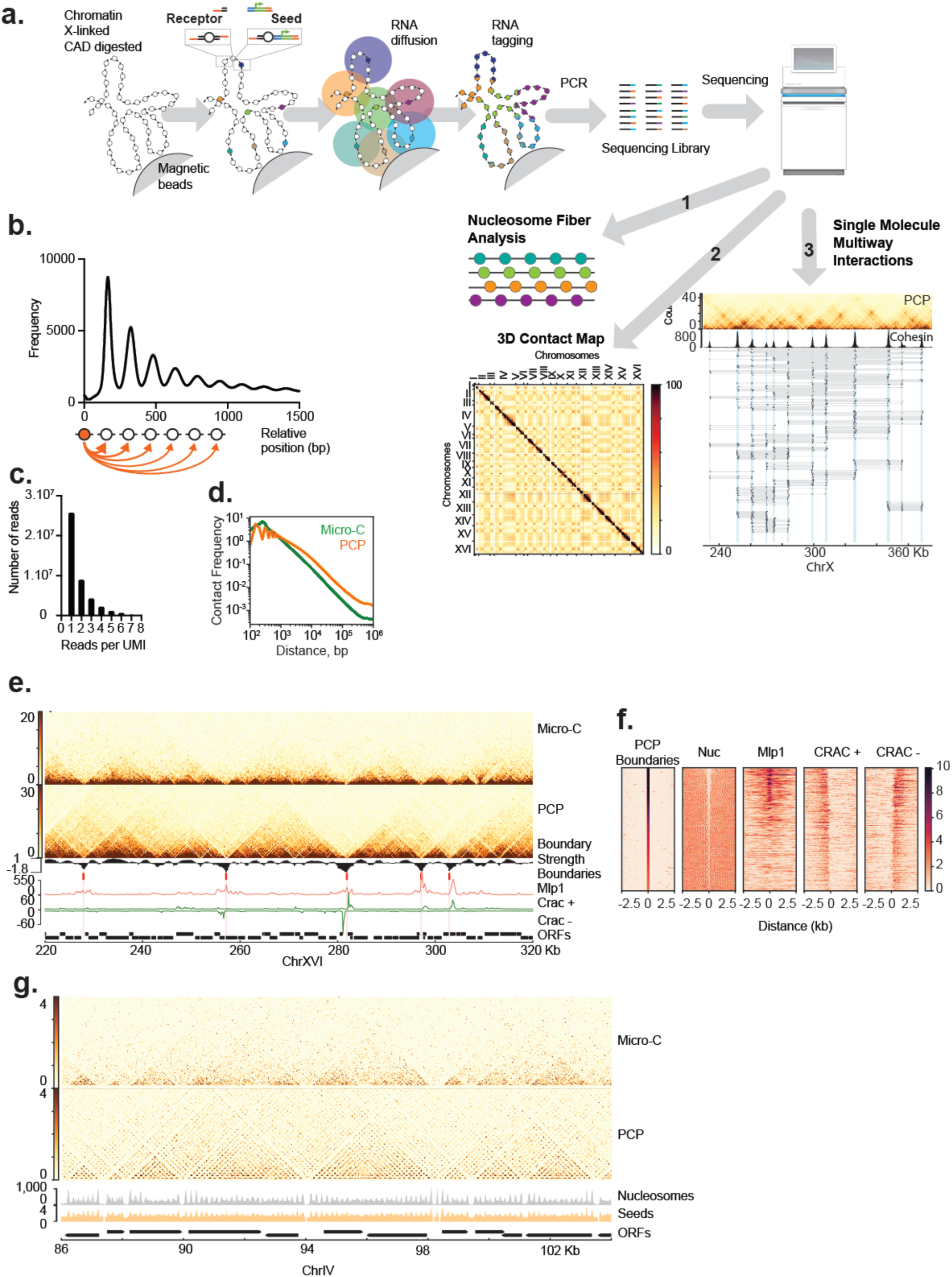
PCP maps 3D genome organization at the nucleosome level. (**a**) Schematic of the PCP reaction on chromatin. Crosslinked chromatin is digested into nucleosomes by Caspase Activated DNAse (CAD) and then immobilized onto magnetic beads. Seed and receptor molecules are ligated into the digested chromatin. The PCP reaction takes place on the beads where RNA tags will diffuse from the seeds and tag nearby receptors. The chromatin is then decrosslinked, the proteins and RNA removed, and a sequencing library generated by PCR. Molecules sharing the same UMI are sorted into tag groups which can be analyzed in several ways. 1. Nucleosome positioning and spacing on individual chromatin fibers. 2. Pairwise interaction maps. 3. Long-range multiway interactions. (**b**) Plot showing the relative frequency of receptor tagging as a function of distance from the seed, the midpoints of the tagged molecules are shown. Seeds that self-tag are excluded from this analysis. (**c**) Plot showing the number of reads per tag group for G1 data. (**d**) Distance decay interaction plots of PCP (orange) and Micro-C (Costantinto et al. 2020, 15 min 23°C sample) (res = 50bp). (**e**) Pairwise interaction landscape obtained from PCP compared to Micro-C (Costantinto et al. 2020) (res = 250bp). PCP detects domains with a higher definition than Micro-C. Strong domain boundaries correlates with Mlp1 ChIP signal (Forey et al. 2020) and transcription activity (Aiello et al. 2020). Insulation score in black, Mlp1 ChIP signal in red, CRAC signal on positive or negative strand in green. (**f**) CID boundaries detected by PCP are sorted by strength. CID boundaries correlate with nucleosome depleted regions (this experiment), transcription (CRAC signal, Aiello et al.2020), and nuclear pores (Mlp1 signal, Forey et al. 2020). (**g**) Nucleosome depleted regions form boundaries of modest strength in the 5’ of genes. Comparison of PCP and Micro-C (Costantino et al. 2020) (res = 25bp). Nucleosome profile derived from the PCP experiment is represented in grey. An even distribution of seeds is observed across the genome and is shown in orange as an example.

Although seeds are ligated into the chromatin at random positions, we can deduce seed location after sequencing which reveals that seeds are evenly distributed across the genome and typically tag receptors within a few hundred base-pairs of their location (Fig 2b, Fig 2g seed location track). We grouped all reads that share the same UMIs into “tag groups” and then calculated all pairwise interactions within each group. Analysis of tag group size reveals that each seed tags a variable number of receptors, which indicates that the PCP reaction has not reached completion, and/or our inability to successfully amplify and sequence tagged molecules (Fig 2c).

Despite some inefficiencies, millions of large tag groups are successfully captured in each reaction. We used the matrix of all pairwise interactions to test if PCP was able to map 3D contacts with similar resolution to Micro-C. Genome-wide, we can observe interactions between the centromeres of the 16 chromosomes, characteristic of the Rabl conformation adopted by the genome of *Saccharomyces cerevisiae* in interphase (Fig S2a) (26, 27). At the chromosome level, PCP can detect long-range interactions as observed between the loci HMR and MATa, characteristic of Mat a strains (28, 29) and low abundance loop regions near centromeres in G1 (Fig S2b). Chromosomal Interaction Domains (CIDs) are detected with increased definition and hierarchy compared to Micro-C (9). PCP is reproducible (Fig S2c) and reveals that the genome is demarcated by large CIDs that typically span ∼10 to 30 kb (median ∼17 kb) and are bounded by regions associated with the nuclear pore (e.g. Mlp1), which are typically highly transcribed (9, 30) (Fig 2e,f,g and Fig S2d). Within the large CIDs, genes appear as micro-domains, separated by small boundaries at their 5’ end (Fig 2e). Mapping nucleosome positions from PCP data, we can confirm that nucleosome depletion and boundaries are correlated (9, 31) with the strongest boundaries of the largest CIDs being wide and depleted of nucleosomes (Fig 2f,g). To test if interfering with the NFR (nucleosome-free region) would erase boundaries, we depleted the catalytic subunit of the chromatin remodeler RSC from the nucleus using an anchor-away system (32). In the absence of Sth1, a subset of NFRs are lost, (32, 33) which results in the loss of the corresponding boundaries at these loci (Fig S3a,b). However, the strongest boundaries, associated with nuclear pores and high transcription are not affected by RSC depletion (Fig S3c).

### High-resolution nucleosome contact maps

High-resolution PCP maps reveal information about individual nucleosomes not available by other methods (Fig.2g), so we investigated how nucleosome positions and contacts change in relation to gene activity. We performed meta-analysis of nucleosomal interactions centered on the dyad of the +1 nucleosome in genes of more than 2kb in length to prevent ambiguities introduced by small genes. Overall, we find that less active genes exhibit well-positioned nucleosome arrays displaying a grid like contact pattern (Fig 3a,c). The 5’ end of transcribed genes also contain phased, positioned nucleosome arrays, yet this signal decays toward the 3’ end where “fuzzy” nucleosomes dominate the raw nucleosome mapping data (Fig 3b) (3). Fuzziness is not a property of the nucleosomes but results from the averaging of the different positions occupied by a nucleosome over a population of cells. Fuzzy nucleosomes constitute a majority of nucleosomes in eukaryotic genomes yet how such nucleosomes contribute to chromatin structure is not clear. Importantly, the contact maps in PCP reveal that nucleosomes in longer genes that appear as fuzzy, have clear spatial relationships with adjacent, fuzzy nucleosomes that collectively form parallel stripes on the interaction matrix (Fig 3d). This reveals that fuzzy nucleosomes can be arranged in organized arrays with uniform nucleosome spacing but delocalized positioning. Moreover, the presence of multiple parallel stripes indicate that the arrays contain multiple nucleosomes (Fig 3e). We broke the PCP data into “tag groups” which represent collections of mapped nucleosomes from individual chromatin fibers (i.e. nucleosomes tagged with the same UMI) and then plotted these relative to genomic coordinate. Such single-molecule maps show the remarkable consistency of organized arrays across the population of cells (Fig 3c,d), and that some genes transition from phased arrays to delocalized arrays (Fig 3d). Delocalized arrays appear to be variable in size and contain up to ∼7 nucleosomes, similar to the delocalized arrays formed in *in vitro* reconstitution experiments (31). Thus, there is an apparent limit to the size of delocalized arrays such that long genes contain multiple adjacent arrays that are not phased with respect to each other (Fig S4a,b). Further, some very long genes alternate between positioned and delocalized arrays (Fig S4c,d).

**Figure 3:**
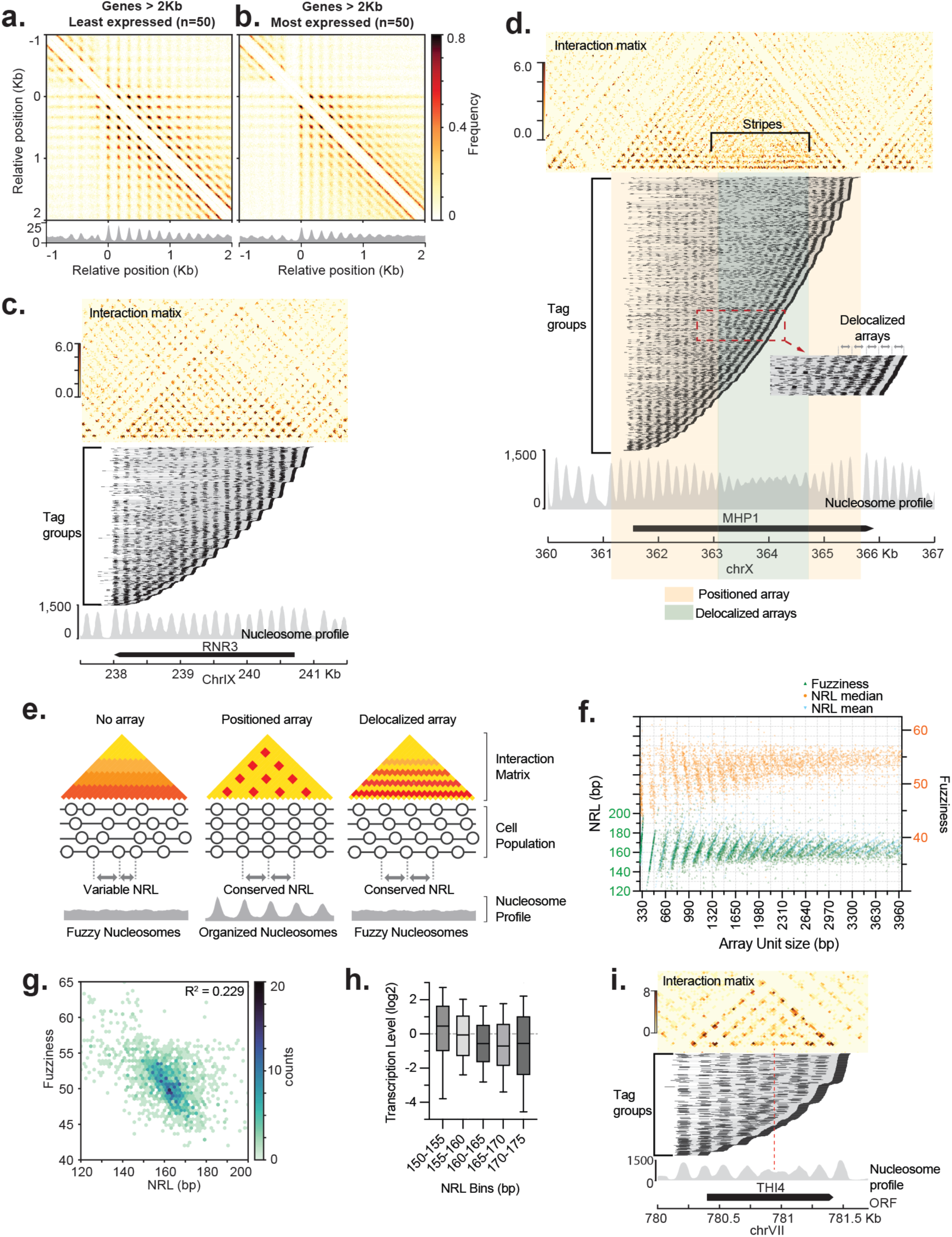
Regularly spaced nucleosome arrays are prevalent over the genome. (**a**) Meta analysis of PCP matrices relative to the dyad of the +1 nucleosome of genes longer than 2Kb, 50 least expressed genes (res = 10bp). (**b**) As **a**, but for the 50 most expressed genes (res = 10bp). (**c**) PCP map of low expressed gene RNR3 (middle, res = 25bp), tag groups are sorted by right most position, nucleosome profile – calculated by read counts per genomic coordinate –is at the bottom in grey. Note the grid-like pattern across the entire gene body. (**d**) PCP map of MHP1 (res = 25bp), tag groups sorted by right most position, nucleosome profile is grey. Note the grid-like pattern at the 5’ end of the gene and parallel stripes at towards the 3’ end of gene in the interaction matrix. (**e**) Schematic of the results observed. No pattern is obtained with poorly spaced, poorly positioned nucleosomes. Dotted pattern is obtained with regularly spaced, well positioned nucleosome arrays. Striped pattern is obtained with regularly spaced delocalized arrays. The number of parallel stripes indicates the number of nucleosomes in the array. (**f**) Scatter plot of median NRL - green dots, mean NRL - blue dots and Fuzziness – orange dots relative to array unit (AU) size. AU are determined by the space between two adjacent NFRs. (**g**) Heatmap of Fuzziness vs NRL for short genes (<2Kb) (**h**) Boxplot of transcription level per bins of NRL length. **(i)** PCP map of THI4 (res = 25bp), tag groups are sorted by right most position, nucleosome profile is grey. Note the heterogeneity within the teg groups showing one or two nucleosomes in the middle of the gene while the nucleosome in 5’ and 3’ remain well positioned.

**Figure 4:**
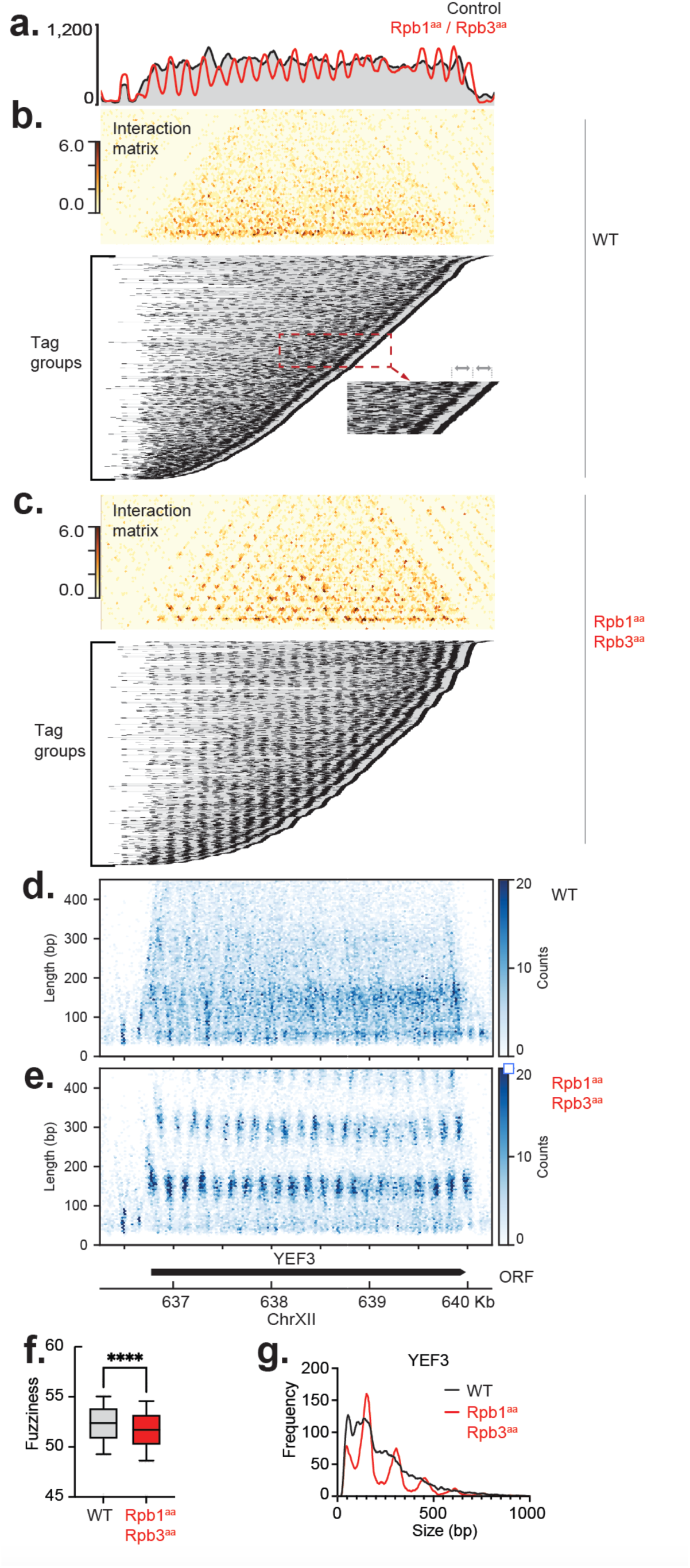
Transcription disrupts chromatin structure. (**a**) Nucleosome profile of YEF3 in WT or when RNA polymerase II is depleted. The profile is calculated by summing the read counts per genomic coordinate (**b**) PCP map of YEF3 in WT while being highly transcribed (res = 25bp), tag groups are sorted by right most position. Note very short, delocalized arrays are found throughout the gene. **(c)** PCP map of YEF3 after RNA polymerase II is depleted by anchor-away (res = 25bp), tag are groups sorted by right most position. Note, ordered nucleosome arrays are now evident across the gene. **(d,e)** Heatmap of read size and abundance over YEF3 in the presence (d) or absence (e) of transcription. The positions and abundance of read midpoints is shown **(f)** Genome-wide fuzziness of AUs in presence (WT) or absence of transcription. **(g)** Read size distribution across the YEF3 gene body in presence (WT) or absence of transcription.

### NFR spacing dictates NRL and Fuzziness

We asked whether features of chromatin architecture influence the prevalence of delocalized arrays. Assuming that nucleosome free regions (NFRs) constrain the extent of nucleosome arrays, we identified all NFRs and then split the genome into array units (AU) which span the distance between sequential NFRs along the genome. We then plotted the median NRL (nucleosome repeat length; calculated as the distance between adjacent nucleosome dyads) for nucleosomes within each AU. As shown in figure (Fig 3f), NRL is related to the length of the AU, consistent with a simple model whereby a distinct number of nucleosomes are packaged within a specific size of AU with upper and lower limits for NRL (34). This pattern is most evident for AUs less than 2kb where the NRL is modulated between ∼155bp to ∼180bp to best accommodate the packaging of nucleosomes between NFRs. Next, we plotted the median fuzziness score for nucleosomes in the same array units. This revealed that fuzziness is typically inversely related to NRL (Fig 3f,g). Thus, the distance between NFRs on either side of genes can generally explain the presence of fuzzy nucleosomes and delocalized arrays: AUs that can accommodate arrays of nucleosomes with an NRL of ∼165-170bp tend to be the least fuzzy are most consistently organized and have the lowest transcription (Fig 3h). AUs too short or too long to accommodate arrays of nucleosomes with a spacing of ∼165-170bp contain increasing numbers of fuzzy nucleosomes. This relationship breaks down when AUs (hence genes) become longer than ∼2 kb as each AU can accommodate different numbers of nucleosomes with minor changes in NRL.

The finding that AUs with lower NRL are less consistently organized is counterintuitive: Tighter packaging should result in reduced variation in nucleosome positions. We examined individual chromatin fibers on a representative short gene (<2 kb) (Fig 3i), which revealed highly heterogeneous chromatin structures where some cells contain tightly packed nucleosomes, whereas others have lower nucleosome occupancy and so can accommodate much wider spacing between the remaining nucleosomes. Thus, the consistency of nucleosome arrays appears to be related to NRL where seemingly tightly packed arrays measured in the population have heterogenous nucleosome positioning and occupancy when viewed with single molecule resolution.

### PCP does not detect 30nm fiber signal nor gene specific organization

The ability of PCP to detect associations between nucleosomes over a wide range of distances led us to investigate whether genes exist in different states of chromatin compaction. Calculating the differences between short range (0.0 – 0.3kb) vs longer (0.3 – 1kb) contacts we find similar trends to those reported with Micro-C (9) where inactive genes have increased longer range contacts, but these effects are generally modest (Fig. S4g). Looking more specifically at longer genes, we compared how interactions decay over a range of distances for genes with different transcription states (Fig.S4f). This analysis showed clear differences in NRL, but minimal changes in interactions, even at longer range. Indeed, similar profiles are observed if we focus on the minority of very highly transcribed genes in the genome such as ribosomal protein genes (Fig. S4g). Further, the smooth decay in proximal inter-nucleosomal interactions indicates that well organized higher order structures, e.g. 30nm fiber, are not a prominent feature of nucleosome arrays detected by PCP (Fig. S4f). While care needs to be taken in generalizing population averaged data such as this, we believe that this data is not consistent with genes being stably maintained in distinct states of chromatin compaction related to their activity. Rather, alterations in chromatin compaction may be transient, local and likely occur with the passage of RNA polymerase II.

### Transcription shut-down affects multiple aspects of chromatin organization

Surveying the genome in G1 arrested cells, it is evident that the most highly transcribed genes have delocalized arrays of nucleosomes across the entire gene. As an example, the *YEF3* gene shows a single stripe on the interaction matrix, indicating that nucleosomes are predominantly organized in pairs (Fig4). This suggests that nucleosome spacing, but not positioning, is rapidly reestablished following the passage of RNA polymerase (RNAPII). To further dissect the effect of transcription, we depleted RNAPII with an anchor away system and asked how this affects chromatin architecture (35). For most genes, depletion of RNAPII has little effect beyond a decrease in fuzziness ((36), Fig4.f, Fig.S5.a-d). But some very highly transcribed genes like YEF3 can be seen to regain organized patterns of nucleosome arrangement, demonstrating that transcription is a cause, rather than a consequence of chromatin disruption (Fig4.a). We then asked how nucleosomes are altered across the gene. Using pair-end PCP data we determined the footprint of mapped reads and found that transcription results in a profound decrease in footprint size across most molecules. Thus, transcription results in a significant gain in subnucleosomal particles – presumably representing partially unwrapped nucleosomes in the process of being transcribed (Fig4d,e,g) (37). Indeed, focusing on the +1 nucleosome we find evidence of progressive unwrapping of the nucleosome in accordance with the direction of transcription which continues to slightly beyond the dyad (Fig.S5.e). We also noticed abundant, sub-nucleosomal signal upstream of YEF3. This signal likely corresponds to a transcription factor. Indeed, Fig. S5.f shows that PCP captures transcription factor footprints, such as that for Rap1, which delimits highly organized and disorganized chromatin at ribosomal protein genes, consistent with Rap1 insulator function (38, 39).

### Mapping multi-way contacts in metaphase chromatin

Comparing G1-with metaphase-arrested cells confirmed a fundamental restructuring of chromatin (27), with clear evidence of cohesin mediated loops (Fig.5a, Fig.S6a) (40–42). PCP interaction maps are similar to those generated by Micro-C, but PCP detects more signal spread throughout Cohesin Associated Regions (CARs) – suggesting frequent interactions between sites of cohesin accumulation and genes with each CAR. We focused on prominent sites of cohesin-mediated loops to ask whether there are structural features of chromatin that may underlie these 3D interactions. We performed meta-analysis of aligned nucleosome arrays at cohesin enrichment sites which revealed a regularly spaced pattern of nucleosomes with an NRL of ∼168bp at the apex of the loops (Fig.5b,c). This spacing is wider than the genome average and is similar to the least expressed genes (Fig.S6b). Of note, the grid-like pattern of nucleosome interactions shown in Fig.5b, does not indicate that interacting regions of chromatin form specific 3D structures, e.g. interdigitated arrays, but does indicate that nucleosome spacing may be one determinant of long-range chromatin interactions.

Chromosome structure is known to be more compact in mitosis and it is assumed that clusters of adjacent cohesin complexes form multi-way contacts, or hubs, generating arrays of adjacent loops (Fig.5d). However, this question is not currently addressable by pairwise ligation methods, such as Micro-C. If cohesin-anchored DNA hubs are formed, multi-way contacts should be evident in the PCP tag groups. Indeed, across chromosomes, we find that tag groups spanning 3 adjacent cohesin peaks are ∼7 fold more abundant than tag groups spanning the same loci in G1, which indicates that multi-way, long range contacts are enriched in mitosis (Fig.5e, Fig.S6c). Further, multi-way contacts are more prevalent across cohesin enrichment sites in mitosis than mock sites, indicating that multiway contacts preferentially form at cohesin enrichment sites, but are not restricted to them. Visualization of individual tag groups that contact adjacent cohesin peaks demonstrates that contacts are formed between a variety of adjacent regions, in addition to cohesin enrichment sites Fig.5f. It is not clear why certain regions are enriched in multi-contact sites, but the variation indicates that a complex and variable mixture of nested loops may be formed. At present, the absolute prevalence of multi-way interactions is not quantifiable, yet the relative abundance of multi-way interactions can be used to measure differences in compaction across the genome. We selected tag groups with ≥3 reads overlapping cohesin peaks and quantitated their abundance across mitotic chromosomes. As shown in Fig.5g, we find that the pericentromeric regions of chromatids display significantly more multi-way cohesin contacts than the chromatid arms, indicating these regions are proportionally more compact. Moreover, the single molecule interaction maps indicate that multiway contacts in the pericentromeric regions appear to have abrupt boundaries (Fig 5g, Fig.S7a,b). This finding is consistent with a compact structure being established at centromeres, distinct from the rest of the genome, and reflecting the unique loading of cohesin at these regions (42–45).

**Figure 5:**
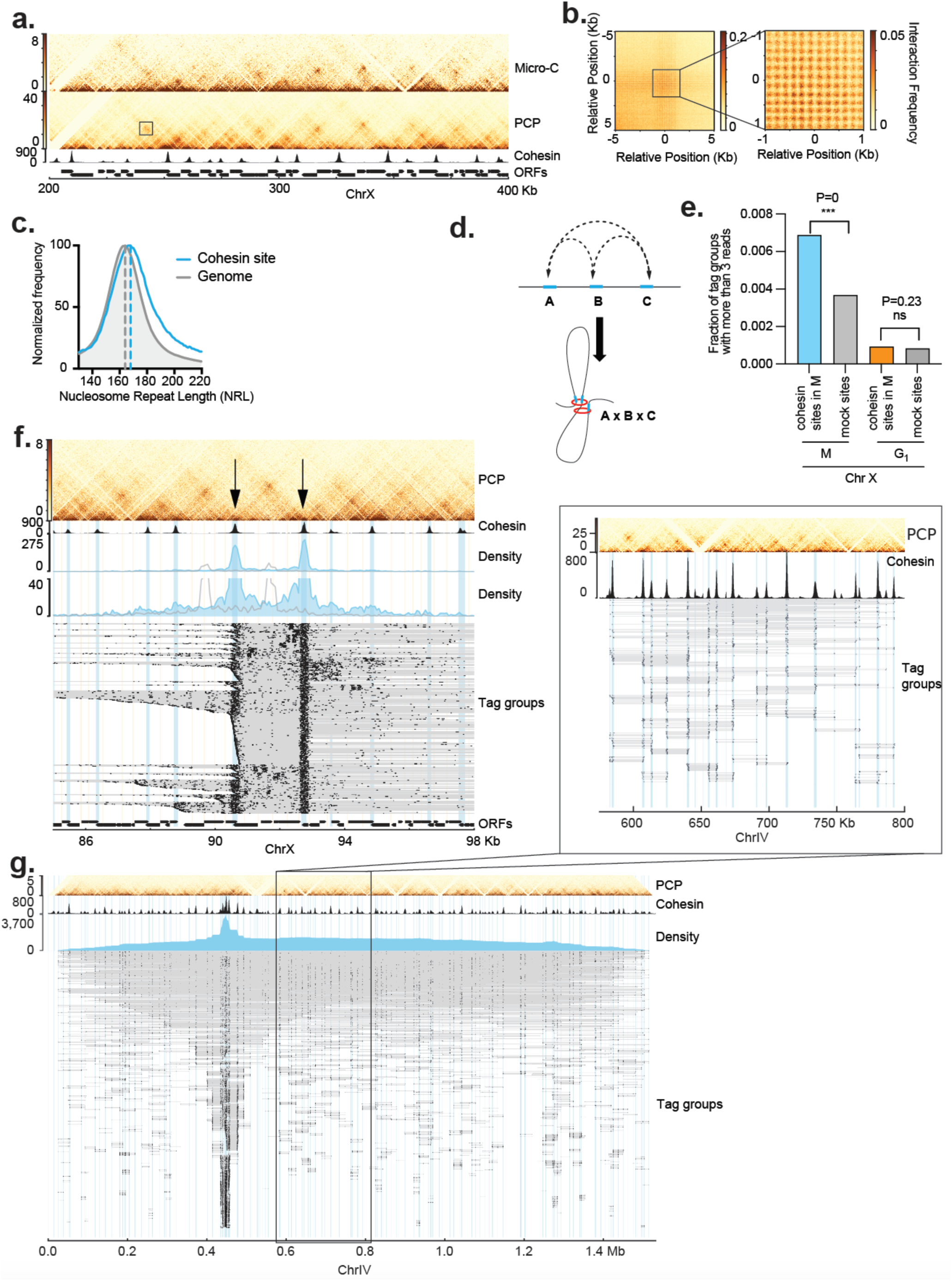
Cohesin organizes yeast chromosomes in multiway interaction hubs in metaphase. (**a**) Comparison of PCP and Micro-C pairwise interaction maps of cells synchronized in Metapahase. Micro-C and Mcd1 (cohesin) ChIP signal is in black (Costantino et al. 2020). (**b**) High-resolution analysis of nucleosome profiles at prominent cohesin interaction sites. (**c**) The nucleosome repeat length at cohesin binding sites is larger (168bp) than the genome average (164bp). (**d**) Schematic of possible multi-way interactions between 3 adjacent cohesin binding sites. (**e**) Fraction of tag groups having reads in at least 3 cohesin binding sites vs mock sites in mitotic chromatin. Comparison is also made for multi-way contacts at the same sites in G1 arrested cells. Fisher exact test is used to calculate p-value. (**f**) Cohesin enrichment sites participate in multi-way long-range interactions. Selecting for tag groups that contain reads in 2 cohesin binding sites reveals complex multi-way interactions. Cohesin ChIP signal is in black (Costantino et al. 2020). Tag groups that contain reads in 2 adjacent cohesin binding sites (black arrows) compared to tag groups that contain reads in 2 adjacent mock sites of same size. Total read coverage is shown in blue for the tag groups overlapping the two cohesin sites or grey for the control. Lower panel is a zoom of the upper intermediate panel. Bottom panel shows each tag group, thick black lines are mapped reads, connectivity of each tag group is shown as grey lines. (**g**) Estimation of relative chromatin compaction levels across chromosomes. Tag groups including at least one read in 3 different cohesin binding sites are mapped along chromosome IV. Cohesin ChIP signal is shown in black (Costantino et al. 2020). Blue area shows the density profile of the reads represented on the lower panel in which each line corresponds to a tag group, thick black lines are mapped reads, connectivity of each tag group is shown as grey lines.

### Overlapping di-nucleosomes form at a subset of divergent promoters

Surveying the genome, we observed intriguing 3D contact patterns within some intergenic regions (Fig.6a). We mapped footprint sizes and observed that the signal is associated with a size of around 260bp, importantly the unusual footprint is of similar abundance to the nucleosome -size footprints nearby, indicating that this entity is a prominent feature of chromatin at this site. Using size as a criteria for stringent selection, we identified ∼60 similar loci, most of which were located between a subset of divergent promoters pairs where the ∼260 bp feature appears to delineate the two nucleosome-depleted regions at the adjacent promoters (Fig. 6b). Comparison with published protein localization datasets (39), and use of motif analysis tools, we were unable to identify any proteins that could account for the unusual footprint (Fig. S8a,b,c). Closer inspection of promoter pairs with a ∼260 bp footprint revealed that the protected regions are flanked by two nucleosome-sized footprints (Fig.6b). Given that the large footprint cannot be explained by a sequence-specific DNA binding factor, we considered the possibility that an alternate nucleosome-like particle may exist at these sites. Evidence now exists to show that overlapping di-nucleosomes (OLDN) can be readily formed *in vitro* through the repositioning of nucleosomes by increased temperature or chromatin remodeling enzymes, such as the yeast RSC complex (46) and mammalian SWI/SNF complexes (47). Indeed, a highly stable OLDN structure has been shown to consist of a histone octamer overlapping a histone hexamer (48). However, there is inconclusive evidence whether OLDNs are a stable, abundant feature of cellular chromatin or whether low abundance footprints of this size are transient intermediates of nucleosome remodeling *in vivo* (49, 50). We reasoned that the population average data shown in Fig.6b indicates that the OLDN is formed by sliding the two adjacent nucleosomes into each other – meaning that the OLDN and adjacent nucleosomes would be mutually exclusive Fig.6c. This hypothesis would be impossible to test with typical population average methods, but using the single-molecule readout of PCP we asked if the OLDN and adjacent nucleosomes are ever found on the same chromatin fiber. Strikingly, selecting for tag groups containing OLDNs we do not find adjacent nucleosomes (Fig.6d). On the other hand, selection for one of the adjacent nucleosomes, we find the other adjacent nucleosome but no reads of a size compatible with an OLDN (Fig.6e). These results show that adjacent nucleosomes and the OLDN are mutually exclusive and are likely being interconverted within the population. If the OLDN is indeed being formed by the remodeling of adjacent nucleosomes, we reasoned that removal of the remodeling enzyme would abolish the OLDN. While depletion of SWI/SNF complex had little effect (not shown), depletion of the catalytic subunit of RSC had three pronounced effects: first, the putative OLDN footprint was significantly reduced in the population and abolished at many individual loci; second, as expected, the relative abundance of the adjacent nucleosomes was drastically increased; third, transcription of the adjacent genes is significantly disrupted (Fig. S6.f,g Fig. S8d-f). Thus, OLDNs appear to be a stable feature of chromatin that is generated by a RSC-mediated remodeling event at specific sites within the genome.

**Figure 6:**
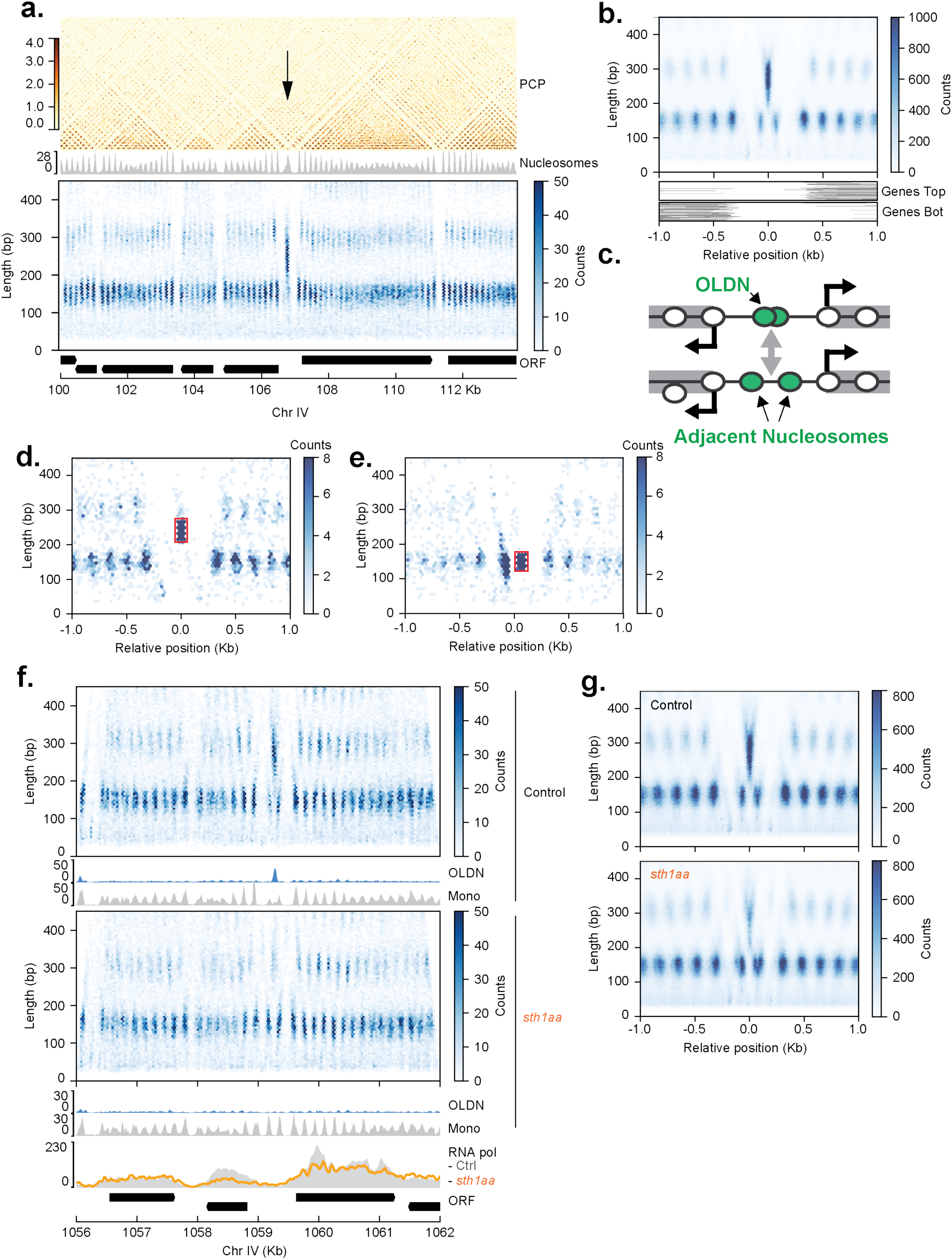
Detection of overlapping di-nucleosomes (OLDN) by PCP. (**a**) PCP maps of a region of chromosome IV with an unusual footprint (black arrow). In grey, nucleosome profile mapped from PCP data. Lower panel represent reads sizes in relation with their positions, color indicates the abundance of the molecule. The divergent gene promoter pair has an unusual footprint that is centered in ∼260bp. (**b**) Metaplot over 61 detected putative OLDN sites in the yeast genome. The unusual molecules footprint is visible in the center of the plot flanked on either side by nucleosomes. Lower plot shows the positions of open reading frames on either the top or bottom strand for this subset of promoters. (**c**) Schematic representing the dynamic interconversion of an OLDN and two adjacent mono-nucleosomes. (**d**) Analysis of single fibers shows OLDN and adjacent nucleosomes are mutually exclusive. Tag groups containing reads that correspond to the putative OLDN were selected, all accompanying reads in the tag groups are plotted to determine which molecules coincide with the OLDN on the same chromatin fiber. Selected molecule size and position is shown in red. No directly adjacent mononucleosomes are detected. **(e)** As (d) except tag groups containing reads that correspond to one of the adjacent mononucleosomes are selected. Selected molecule size and position is shown in red. Note, an adjacent nucleosome is detected at high frequency but there is no signal for the putative OLDN. **(f)** Comparison of footprint positions and size distribution in control (top panel) and RSC depletion (*sth1aa*, lower panel) at a single representative locus. Nucleosome tracks of OLDN size range are blue (210 bp < insert size < 270 bp) and mono-nucleosomes in grey. RNA polII ChIP in both conditions (Kubik et al. 2018) is shown in the bottom panel. (**g**)Metaplot at the 61 sites showing footprint positions and size distribution in control (top panel) and RSC depleted condition (lower panel, *sth1aa*).

## Discussion

We show that a system using diffusible RNA tags can be used as a proximity labeling approach to map multiplex molecular interactions genome-wide and with single-molecule precision. The approach involves the transfer of sequence information from seeds to receptors. The reaction is biased toward tagging receptors near the seed, which allows PCP to be used to map spatial relationships between molecules. PCP differs from conventional proximity tagging approaches as the use of nucleic acids permits a vast library of unique tags to be rapidly produced. Such diversity allows for multiplex reactions to be conducted with the number of unique tags dictated by the size of the UMI on the seed. The PCP assay has several tunable parameters and further optimization will likely improve aspects of the method. Increasing the local tagging efficiency should allow more detailed single-molecule analysis of nucleosome positioning on individual chromatin fibers. In addition, PCP can be tailored to specific needs. For example, altering the seed:receptor ratio should allow interactions to be assessed at different spatial resolutions. Seeds or receptors could also be targeted to specific genomic locations or attached to specific proteins, which would allow targeted analysis of regions of interest, or the occupancy of protein(s) or RNA to be integrated into the high-resolution 3D contact maps. For chromatin analysis, we found that CAD is superior to MNase. Principally, this is because CAD is far less likely to over-digest nucleosomal DNA. Thus, optimization of CAD digestion is trivial and essentially all DNA ends resulting from digestion can be repaired and ligated, which is not the case with MNase (Fig S1.b). Such efficiency is essential for single-molecule assays such as PCP that have the ability to map most molecules (e.g. nucleosomes) put into the reaction. Given this sensitivity, very low input amounts of chromatin are required. Importantly, this will likely allow single cell analysis of metazoan genomes to be conducted with few modifications.

By mapping the local interactions of nucleosomes, we show that nucleosome arrays dominate the chromatin landscape. This finding is consistent with the classical nucleosome ladder generated by digestion of cellular chromatin with MNase (51, 52) and long-read genomics methods to map nucleosome arrays (18, 53). Yet, because PCP maps the positions and connectivity of individual nucleosomes, we derive a more complete understanding of nucleosome positions, array formation and longer-range contacts. The presence of delocalized arrays is more prevalent in transcribed genes, yet transcription cannot explain all delocalized arrays. We provide evidence that fuzzy nucleosomes and delocalized arrays can be explained by a simple packaging principle governed by the distance between NFRs at the ends of genes. All genes (with the exception of the most highly transcribed) have 2-5 positioned nucleosomes abutting the NFR at their 5’ and 3’ ends (1). The locations of nucleosomes in the middles of genes are controlled by nucleosome spacing factors which space nucleosomes relative to each other, and the positioned nucleosomes at the gene ends. NRL is related to fuzziness, and genes that can accommodate an array of nucleosomes with an NRL of ∼165-170bp between NFRs typically contain organized arrays that are phased with respect to the positioned nucleosomes at both gene ends. Such genes tend to be the least expressed genes in the genome. However, as the distance between NFRs at gene ends varies, nucleosomes in the middle of the array must adopt an average NRL either shorter or longer than ∼165-170bp (Fig.3f). Deviation in NRL appears to generate less stable arrays with heterogenous nucleosome occupancy (Fig.3I). As genes become longer, an increasing number of nucleosomes in the middle of genes are controlled by spacing factors so they are in register with nucleosomes at one of the two ends of the genes. Spacing factors act to regulate the amount of linker DNA between nucleosomes, and we find that delocalized arrays contain up to ∼7 nucleosomes. Yet repositioning a nucleosome array in one direction will create longer stretches of free DNA in the other – favoring subsequent repositioning towards the available DNA. This dynamic competition ultimately gives rise to the large, delocalized nucleosome arrays we observe in PCP.

We find little evidence that genes stably exist in different states of chromatin compaction and conclude that subtle differences in compaction are transient and a consequence of ongoing transcription, much like the patterns of histone modifications seen in gene bodies (54). Measured connectivity between adjacent nucleosomes decays uniformly with distance and we find no evidence of distinct higher order folding patterns expected for 30nm fibers, while this finding may reflect that PCP is insensitive to such structures, our data is consistent with an increasing body of evidence that show highly-structured 30nm fibers are unlikely to exist *in vivo* (55–57). Nucleosome repeat length does appear to relate to stability of nucleosome arrays. Indeed, arrays with a consistent NRL of ∼165-170 bp have the most consistently positioned nucleosomes. This spacing may favor more stable interactions between adjacent nucleosomes (58). Nucleosome spacing patterns are likely interpreted by remodeling/modifying factors. For example, Rpd3, a histone deacetylase, associated with gene repression preferentially deacetylates nucleosome arrays with a longer NRL (59).

The novelty of PCP compared to proximity ligation assays lies in its ability to map multiway long-range interactions. Recent studies using Hi-C and Micro-C have revealed the formation of cohesin-dependent loops in G2 phase. These interactions between consecutive cohesin binding sites led to a model in which consecutive loops aggregate, potentially forming hubs of cohesins. However, the pairwise nature of interactions detected by Micro-C and Hi-C has limited the ability to directly test this model, leaving open the possibility that cohesin may enforce pairwise interactions while excluding multiway clustering. Here, we provide direct evidence that cohesin forms multiway cohesin hubs across the genome which provides a novel measure of chromosomal compaction. Furthermore, the increased resolution afforded by PCP reveals that sites of cohesin accumulation coincide with regions exhibiting wider nucleosome repeat lengths similar to repressed genes. By enabling the simultaneous detection of nucleosome organization and long-range interactions on the same molecule, PCP offers a powerful approach to investigating the interplay between chromatin architecture and SMC complexes. This dual capability promises to shed new light on the mechanisms governing genome organization and stability.

The precise mapping of DNA footprints resultant from CAD digestion of chromatin has allowed us to generate evidence that alternate nucleosome structures exist *in vivo*. In this case, we show that an OLDN is specifically formed by the activity of the RSC chromatin remodeling enzyme at a position that separates two adjacent gene promoters; importantly, foot-printing data shows that the OLDN is as abundant as the adjacent nucleosomes. The single-molecule readout of PCP allowed us to determine that the OLDN is being formed from two adjacent nucleosomes in a manner that appears to regulate accessibility of the two gene promoters. The OLDN particles we describe appear to be an abundant, relatively stable component of chromatin, and indicate that such particles are generated for specific purposes. The highly defined positioning of OLDNs suggests they may act as a barrier or insulator separating closely spaced promoters. An intriguing possibility is that the unusual structure of the OLDN may be refractory to certain chromatin remodeling enzymes or chaperones – such as FACT – which bind to specific areas on nucleosomal structures (60). Thus, OLDNs may pose an obstacle to the transcription machinery to help enforce the directionality of transcription. Given the highly conserved nature of chromatin and chromatin remodeling enzymes, similar, stable OLDNs are likely generated across eukaryotes.

In sum, we demonstrate that PCP can be used to map chromatin with high resolution while also capturing long rage contacts. The simultaneous mapping of transcription factors, nucleosomes, and their arrays as well as multiplex 3D genome folding has allowed us to provide new insights into fundamental aspects of chromatin biology. The breadth of data produced from PCP should ultimately allow more wholistic understanding of many chromatin transactions that occur in cells.

### Limitations

A limitation of the PCP approach is the large number of sequencing reads required to profile large genomes with single nucleosome resolution, yet this reflects the vast complexity of genomes and will limit any single molecule approach. While PCP is significantly more efficient than 3-C based methods, not all receptors are tagged in each assay and so single-fiber analysis is currently sub-optimal. Furthermore, single-fiber analysis is complicated by the inability to discern whether missing data (e.g. a nucleosome) is a reflection of biology or a detection failure. For PCP to work effectively, a competition between nearby seeds in chromatin should be established to ensure that local tagging is favored over long-range non-specific tagging. We assume that all seeds in the reaction are equally competent to generate RNA tags capable of annealing to receptors and being copied by reverse transcriptase. Incompetent seeds/RNA tags would diminish the tagging efficiency and potentially increase non-specific tagging. We fail to detect distinct nucleosome arrangements consistent with a 30nm fiber or large changes in chromatin “compaction” when comparing expressed vs non expressed genes. While our conclusions are consistent with a growing body of evidence, deficiencies in PCP may underlie our negative data.

## Acknowledgments

We thank Slawomir Kubik and David Shore, as well as Thomas Guerin and Frank Uhlmann for sharing yeast strains. We thank Tiffany Willis for help with yeast strain growth and Toshio Tsukiyama for the critical reading of the manuscript. We thank members of the Whitehouse and Remus lab for sharing reagents and materials.

## Author contributions

Conceptualization: IW, AD

Experiments: AD, BB, JY

Sequencing: NM

Data analysis: AD, IW, RK

Funding acquisition: IW

Writing: IW, AD

## Data and materials availability

Code, raw and processed data will be available on GEO and SRA after acceptation of the manuscript.

## Supplementary Materials

### Materials and Methods

#### Cultures for PCP

75mL of 1.10^7^ cells.ml were grown in YPD and fixed in 1% formaldehyde for 15min at room temperature (RT). Formaldehyde was quenched with 0.75M Tris HCl, pH 8 for 1min. Cells were washed three times in 1X PBS at 4°C. Cells were resuspended and flash frozen in 300µL of XLB3 (Sorbitol 1M, NaCl 50mM, HEPES pH 7.5 10mM, MgCl2 1mM). For culture in G1, cells were synchronized with ⍺-factor at 2µg.mL for 3hr with re-addition of ⍺-factor every hour before fixation. For culture in Metaphase, cells were synchronized in G1 as above, released by filtration into YPD containing Nocodazole 15µg/mL (sigma, M1404) for 2h before fixation. For anchor away depletion, Rapamycin (sigma, 553210) was added to the media 1hr before fixation at 1µg.mL. For Auxin Induced Degradation, IAA (sigma, I3750) was added at 0.5mM. The strains used can be found in the Supplemental Procedures section.

#### CAD purification and activation

pETDuet_huCAD_ICADL (Addgene, 100098) was transformed into T7 Express LysY/Iq competent cells (NEB, C3013I). Cells were grown in 2.4L of LB2YT (Tryptone 10 g/L, Yeast Extract 10 g/L, NaCl 5 g/L) until OD reached 0.38 and induced with IPTG 0.5mM for 4h at 37°C. Cells were pelleted at 4000 x g for 30min and the pellet was stored at - 80°C. The pellet was resuspended in 70 mL Lysis buffer (NaCl 300 mM, HEPES pH 7.5 50 mM, Glycerol 10% w/v, Imidazole 20 mM, 1X Protease Inhibitor (Pierce, A32963) and sonicated on ice for 10 seconds on, 20 seconds off at 50% amplitude for 6 min. The sonicated mix is transferred to 2 x 40 mL high speed centrifuge tubes, and spun down at 18,000 x g for 1 hour at 4°C. The supernatant was recovered and passed through a HisTrap FF (Cytvia, 17528601). The column was washed with 25 mL of Wash Buffer (NaCl 300 mM, HEPES pH 7.5 50 mM, Glycerol 10% w/v, Imidazole 40 mM, 1 x Pierce Protease Inhibitor Cocktail Tablet). The complex was eluted with 6mL of Elution Buffer (NaCl 0.3M, HEPES pH 7.5 50 mM, Glycerol, 10% w/v, Imidazole 0.5 M) and stored at -80°C. The purified CAD/ICAD was dialyzed into Exchange Buffer (NaCl 400 mM, HEPES pH 7.5 20 mM, Glycerol 10% w/v, DTT 5mM) and aliquoted in 100µL of 50µg of protein. On the day of the experiment, 87 µL of CAD was activated by addition of 10µL of TEV buffer and 3µL of TEV enzyme for 1hr at 30°C (NEB, P8112S).

#### Linkers annealing and seed production

Seed linkers, and PCP linkers, were independently mixed at 25µM in Annealing Buffer (100 mM KAc; 30 mM HEPES pH 7.5). Annealing was performed by heating to 94°C for 4min, and letting the heat block cool down to RT overnight. Before ligation to chromatin, Seed linker and PCP linker were mixed to the desired ratio, in this study 1:10 seedlinker to PCP linker ratio. Seed were produced as follows: 1 ng of the seedbase sequence was amplified by PCR with the oligos and with Terra (Takara, 639270) for 15 cycles. The PCR reaction was supplemented with 2µL of dNTP and 0.5µL of Klenow fragment (NEB, M0210) for 30min at 20°C to repair the UMI. The product was purified through Zymo columns (Zymo research, D4014) and eluted in 10µL, the ends of the molecules were blocked with ddGTP using Terminal Transferase (NEB, M0315) following manufacturer’s instruction. The product was purified again with Zymo columns and digested ON at 37°C with BcoDI (NEB, R0542). The digested product was resolved on a 1.8% agarose gel and the seed was gel purified (∼ 130bp product). Seed was diluted to 10ng.µL and stored at 4°C.

#### PCP on chromatin

Cells were broken using Freezer Mill (Spex SamplePrep, 6875D) for 5 cycles, 2 min, 15cps. After verification of the breakage under the microscope, cells were resuspended in 450µL of XLB3 (supplemented with Protease inhibitors, A32955, Pierce) and cross-linked by addition of DSG at 6mM final (Disuccinimidyl glutarate, A35392, ThermoFisher) for 30min at 30°C. DSG was quenched by addition of 50µL of 1M Tris HCl, pH 8 for 1min. Cells where wash 3 times HS buffer (Sorbitol 1M, Hepes pH 7.5 10mM) and resuspended in CAD buffer (Sorbitol 1M, Hepes pH 7.5 10mM, DTT 5mM, IGEPAL 0.1%, MgCl2 1mM, Protease inhibitors A32955, Pierce) and supplemented with pre-activated CAD. Chromatin was digested for 1hr at 37°C with 1000 rpm agitation. Cells were placed on ice for 2 min and washed twice in SB buffer (10mM Hepes pH 7.5, 0.1% IGEPAL, 1mM EDTA, 1mM DTT) and resuspended in 300µL of SB. 100µL of cells were diluted in 300µL of SB and added to 50µL of pre-activated NHS beads (Pierce, 88827) and agitated for 10min at RT. Bead binding was quenched by addition of 30µL of 1M Tris HCl for 1 min. Beads were resuspended in 500µL of SB and supplemented with 3mM ZnCl2 final for 2min. Beads were washed twice in SB and resuspended in 500µL of SB. End-Repair and A-tailing was performed with the NEBNext Ultra II DNA library Prep Kit (NEB, E7645) for 30min at 20°C with constant mixing, then 30min 65°C with constant mixing. Cells were incubated for 2 min on ice and washed twice in HB (10mM Hepes pH7.5, 1M NaCl, 0.5% Na-deoxycholate, 0.5% Triton X-100, 0.5% IGEPAL) and twice in SB buffer. Beads were resuspended in 100 µL of Hepes pH 7.5 10mM. 20µL of beads were incubated with 2µL of linker mixture at desired ratio at 25µM and 22µL Blunt/TA ligase (NEB, M0367L) for 40min at 20°C. Beads were washed twice in HB and twice in SB. Beads were resuspended in 20µL of Hepes pH 7.5 10mM with 2µL of seed at 10ng.µL and 22µL of Blunt/TA ligase (NEB, M0367L) for 40min at 20°C. Beads were washed twice in HB and twice in SB and resuspended in 20µL of Hepes pH 7.5 10mM.

PCP reaction were conducted on 1µL and 0.25µL of beads mixture in PCP buffer (HEPES pH 7.5 40mM, 6mM MgCl2, 0.5mM dNTP, 0.5 mM rMTP, 5mM DTT, 2.5% PEG 20K, 0.5 µL RNase inhibitor (Pierce, AM2694), 1µL of Reverse Transcriptase (ProtoScript II, NEB M0368), 3µL of T7 RNA Polymerase (NEB, M0251)) for 1hr at 30°C. PCP reaction was stopped with 20µL 100mM Tris HCl pH 7.5 with 0.1 µL of RNAse cocktail (Invitrogen, AM2286) for at least 2 hours at 65°C. The decrosslinked chromatin was supplemented with 17 µL of H2O, 1.5 µL of each of the PCR primers (any of the Illumina NEBNext primers, E6440S) at 10 µM and 50µL of KAPA RM (Roche, 07958927001) and incubated in the PCR reaction for 12 cycles (5min 95°C, [20 sec 98°C, 15 sec 71°C, 1min 72°C] x12, 2 min 72°C, 4°C). DNA was purified using Zymo columns (Zymo research, D4014) and resuspended in 10µL. If the amount of DNA recovered is close to 50ng (± 5ng), the library was submitted for sequencing. Otherwise, PCP reaction was restarted with adjustment of the input.

#### Data processing

Details of data processing will be provided in supplemental procedures and on GitHub (https://github.com/Axel-Delamarre/Whitehouse-Lab/). In principle, UMIs are extracted with UMI-tools (61). Only the reads containing the linker sequence are analyzed. Reads containing the seed linker sequence were separated and trimmed and aligned separately. After alignment with Bowtie2, Seed and Acceptor molecules BAM files were merged, sorted, then deduplicated using UMI-tools. Reads sharing the same UMI and whose midpoint is separated by less than 10bp were excluded, they correspond to the different strands of the same molecule. Reads were grouped based on their UMI sequences. For Matrix production, pairwise interactions were deduced from the UMI groups and matrix were produced with Cooler and Juicer packages. Boundaries were called using cooltools (62), bin of 250bp and window of 1.5kb. The matrices are not balanced (cf. supplemental procedures). Reproducibility was measured by pearson correlation with hicexplorer hiccorrelate between replicates (≥0.9).

### Supplementary Text

#### Matrix Balancing

Matrix balancing is a common practice in analyzing 3C (chromosome conformation capture) results to normalize data, ensuring that each row and column sum to the same value. This approach helps to mitigate genome coverage and digestion biases, providing a more uniform representation of each restriction fragment. However, in technologies like Micro-C and PCP, which essentially map nucleosomes, the natural biological variations—such as nucleosome-free regions (NFRs) that typically have lower signals—become significant. Balancing in these contexts can introduce noise by reducing the signal on nucleosome dyads and increasing it on linkers and NFR. Consequently, the matrices in this article are not balanced.

#### Strains

yIW385: CEN.PK -- MATa -- URA3; TRP1; LEU2; HIS3; MAL2-8C; SUC2

yIW709: YSK111 -- W303 -- tor1-1 fpr1::NAT RPL13A-2XFKB12::TRP1 STH1-FRB-KanMX; matalpha

yIW711: HHY190 -- W303 -- tor1-1 fpr1::NAT RPL13A-2XFKB12::TRP1 SNF2-FRB-KanMX; matalpha

---

**Fig. S1.**
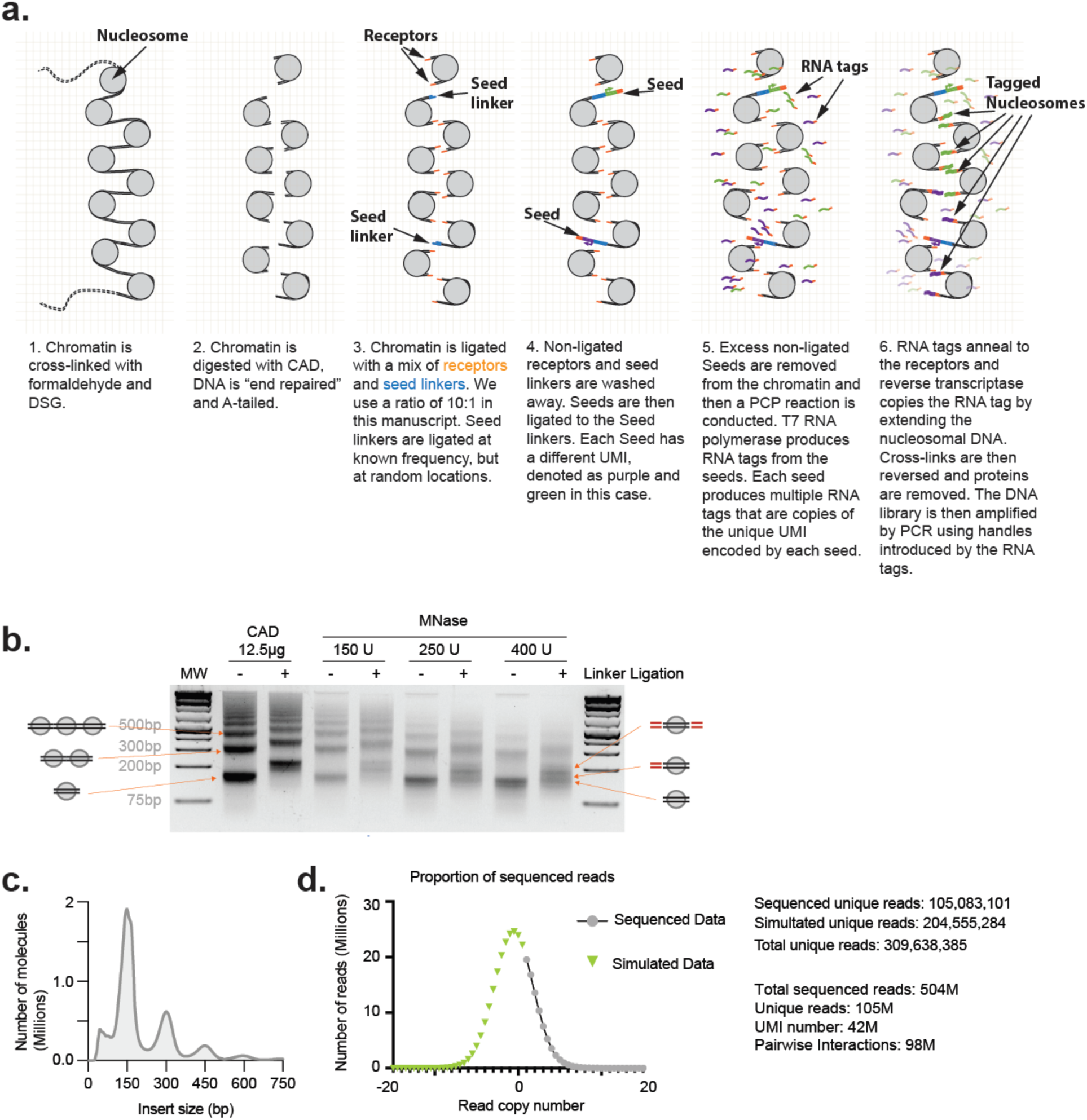
a. Detailed description of PCP on chromatin. b. Ligation efficiency of short DNA linkers to nucleosomal DNA digested by MNase or CAD in crosslinked chromatin. Following ligation, crosslinks and proteins are removed and the purified DNA is migrated on a 1.8% agarose gel. c. Insert size distribution of the PCP library. d. Basic statistics for the G1 PCP sequencing experiment. Graph shows number of times each read was sequenced and estimation (by gaussian curve fit) of the total amount of unique sequences present in the library (green). From this we assume that 30% of the diversity of the library was sequenced.

**Fig. S2.**
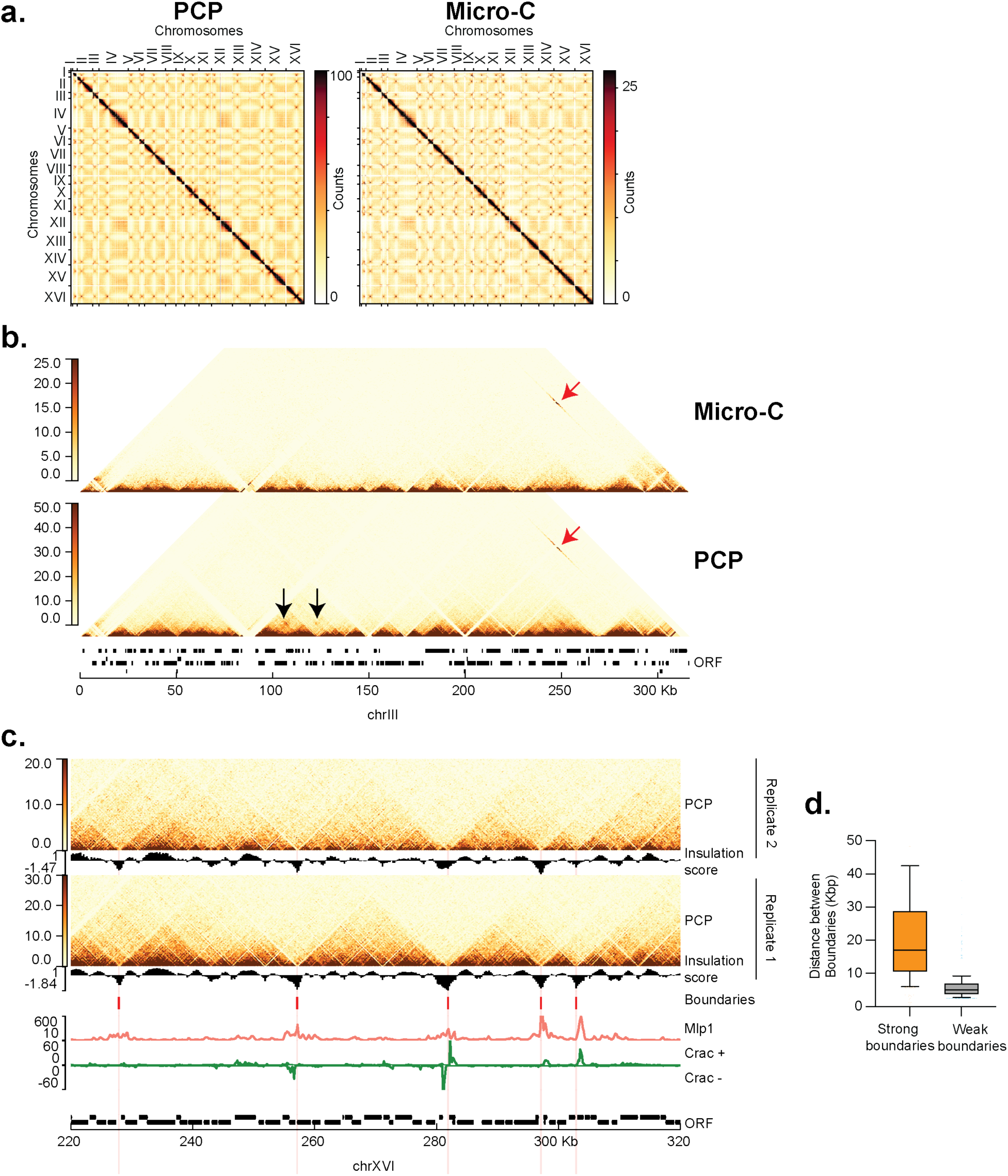
a. Comparison of whole genome pairwise interaction maps of PCP and Micro-C (40) for all chromosomes showing expected interactions of centromeres as dots on the matrix. b. Matrix of pairwise interactions over chromosome III of PCP and Micro-C (40) (res = 500bp). Long-range interactions between the HMR and Mat**a** locus are detected (red arrow) as well as novel loops close to the centromere in the PCP data (black arrows). c. Comparison of 2 replicates of the PCP experiment in G1. (res = 250bp). Replicate 1, 105M unique reads. Replicate 2, 67M unique reads. Pearson correlation = 0.95. d. Distance between strong and weak CID boundaries. Boundaries were called with cooltools (62) at resolution 250bp, with a window of 1.5Kb. Strong boundaries > 1.75, weak boundaries > 0.5. Median, Box and Whiskers 10-90 percentiles.

**Fig. S3.**
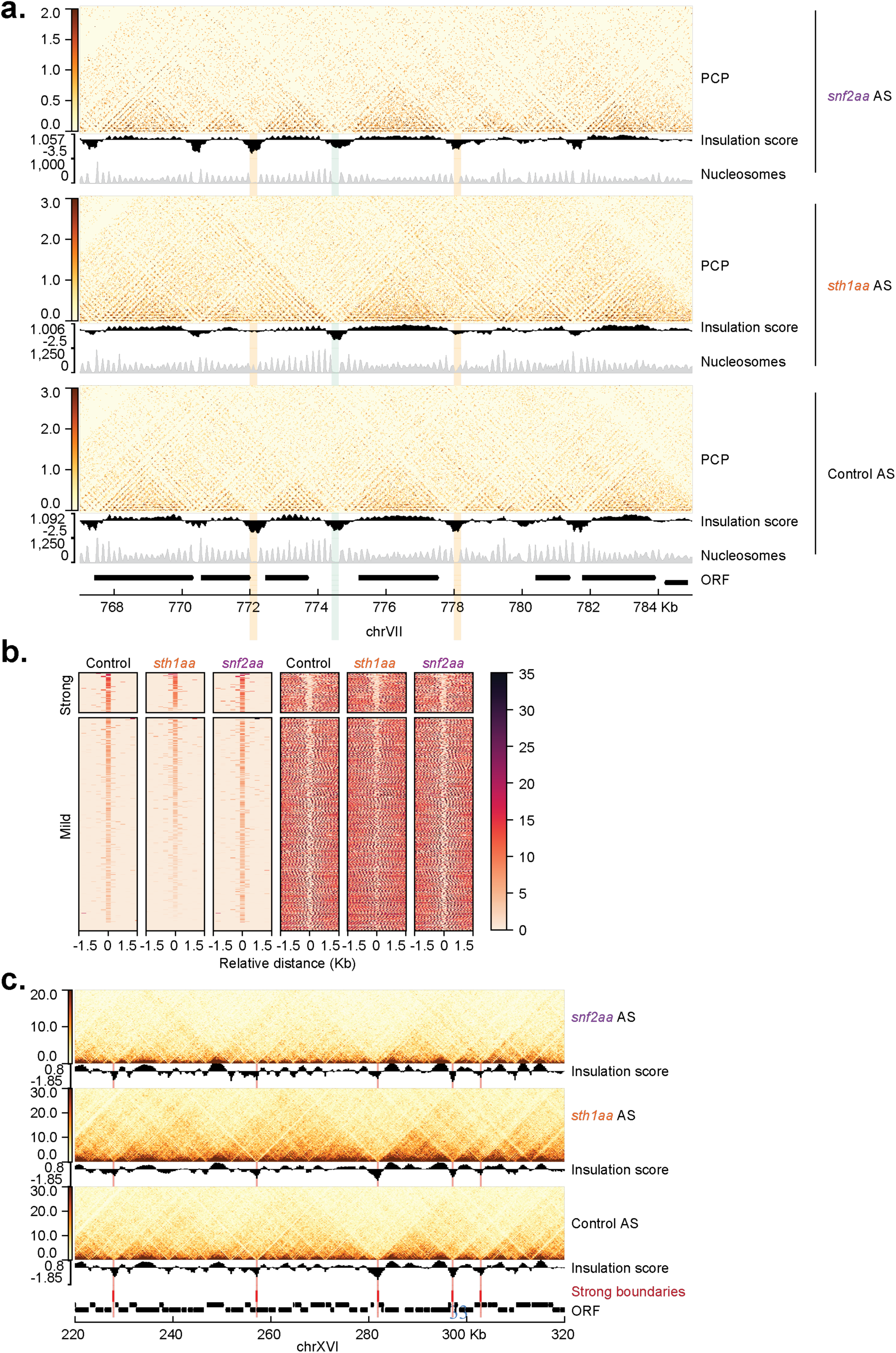
a. Pairwise interaction landscape of asynchronously (AS) grown culture of control, RSC depleted (*sth1aa*) and SWI/SNF depleted (*snf2aa*). Insulation score is in black. Strong domain boundaries (red) are not affected by chromatin remodeler depletion. (res = 250bp). b. Pairwise interaction landscape of asynchronously grown culture of control, RSC depleted (*sth1aa*) and SWI/SNF depleted (*snf2aa*). Insulation score is in black. Weak boundaries at which the nucleosome positioning is affected by RSC depletion are affected (Orange highlights). Weak boundaries at which the nucleosome positioning is not affected by RSC depletion are not affected (Green highlight). (res = 25bp). c. CID boundaries detected by PCP are sorted by strength. Nucleosome profile alteration correlates with CID strength alteration in RSC depletion (*sth1aa*).

**Fig. S4.**
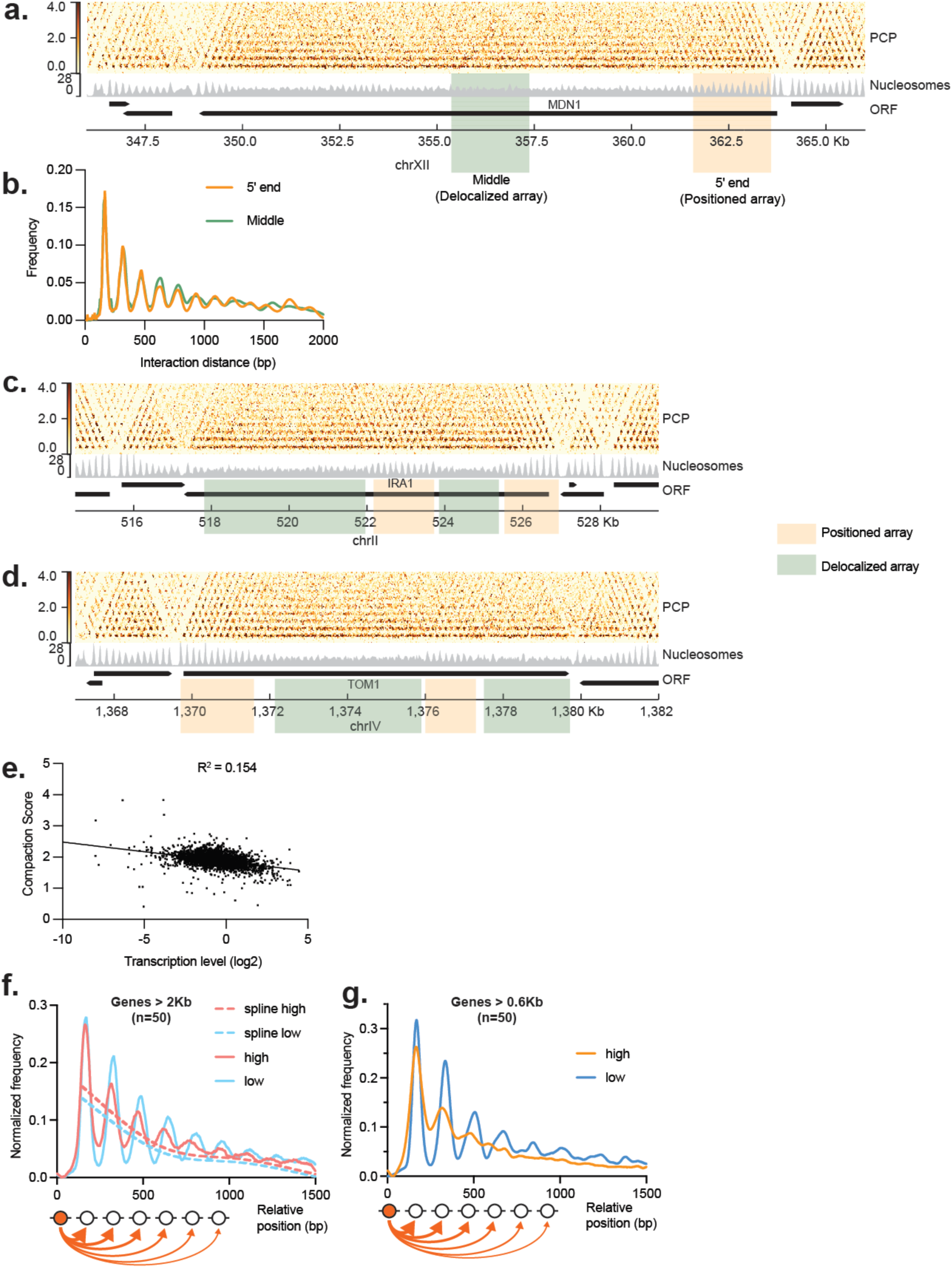
a. PCP map of the long gene MDN1 (res = 25bp), Nucleosome profile (grey). Note the grid-like pattern at the 5’ end of the gene and parallel stripes towards the middle and 3’ end of gene. b. Interaction distance decay profile over 2kb at the 5’ end of MDN1 (orange highlight in a. and line in b.) and 2kb in the middle of MDN1 (green highlight in a. and line in b.). Despite the difference in pattern the NRL is similar between these two regions. c. PCP map of the IRA1 gene (res = 25bp), Nucleosome profile (grey). Note the variable locations of the positioned and delocalized arrays. d. As **c** for the TOM1 gene. e. Scatter plot of transcription level vs compaction level. Transcription data, Aiello et al. 2020. Compaction was calculated as the ratio of longer (0.3-1Kb) over shorter (0-0.3Kb) interaction frequency as proposed earlier (9). f. Frequency of tagging relative to the seed position for the subset of genes used in a and b. The most expressed genes are in red and least expressed genes in blue. g. Frequency of tagging relative to the seed position for gene of at least 600bp. The most expressed are in orange and the least expressed is in blue.

**Fig. S5.**
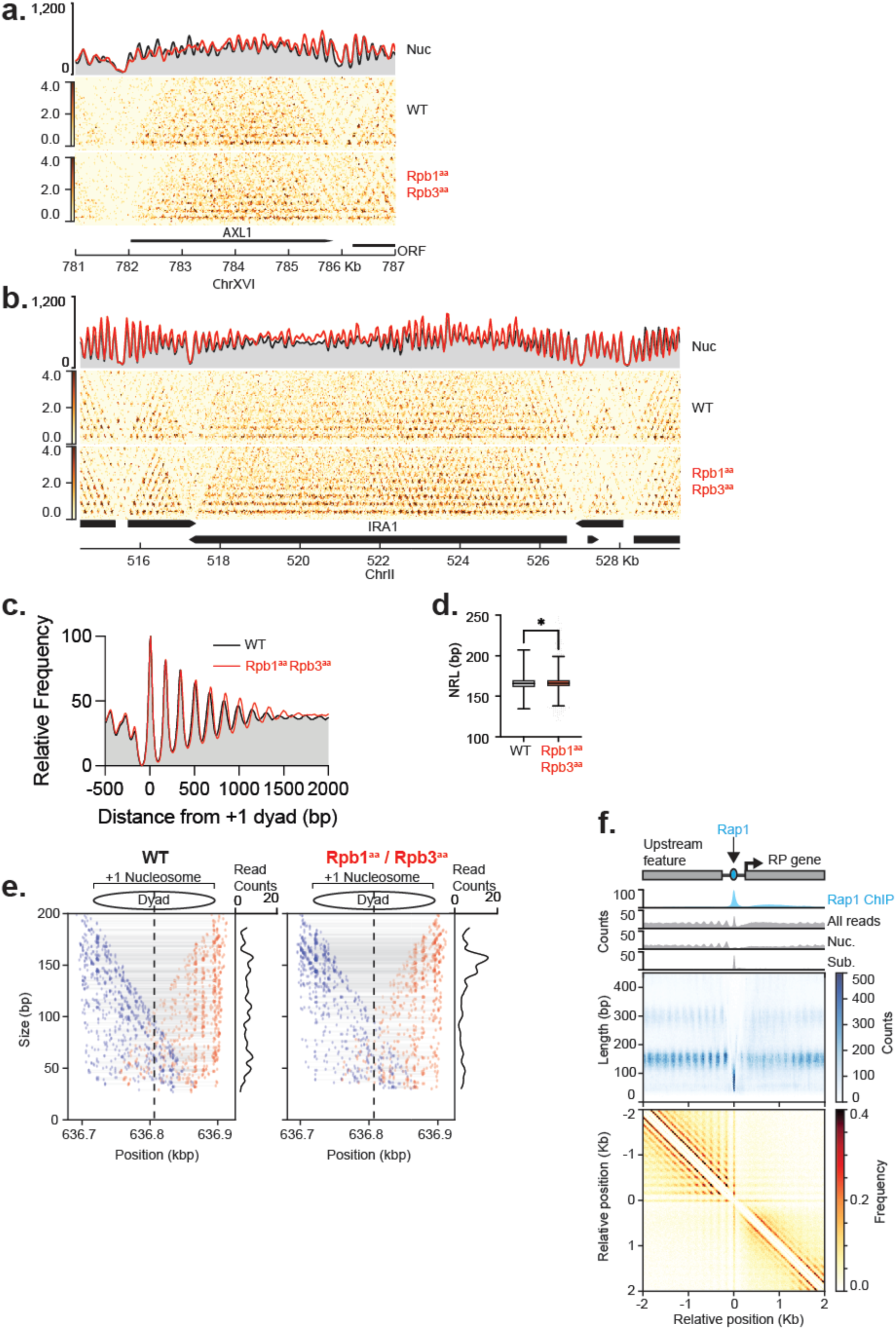
a. Nucleosome and PCP map of gene AXL1 before (grey) and after RNA polymerase II is depleted by anchor-away (red). (res = 25bp) b. Nucleosome and PCP map of gene IRA1 before (grey) and after RNA polymerase II is depleted by anchor-away (red). (res = 25bp) c. Average profile of nucleosome profiles centered on the dyad of the +1 nucleosomes in the presence (black) or absence (red) of transcription. d. Genome-wide Fuzziness of AU in presence or absence of transcription. e. Plot of each read of the +1 nucleosome of YEF3 in the presence (left panel) or absence (right panel) of transcription. Blue dots correspond to the 5’ end of the read, red dots correspond to the 3’ end of the read, grey bar represent the read. f. Meta plot of highly transcribed, ribosomal protein genes. The genes are organized with respect to the Rap1 protein at position 0. Top panel, Rap1 ChIP-MNase in blue (Gutin et al. 2018), All reads size, Mono-nucleosomal (Nuc.) reads only and Sub-nucleosomal reads only (Sub. 20 bp < insert size < 90 bp) are represented. Medium panel represent the density of read by position relative to insert size from mapped PCP data. The bottom panel is a pile-up analysis of PCP matrix oriented relative to the RP genes (res = 10bp). Note that the gene body contains short, delocalized nucleosome arrays.

**Fig. S6.**
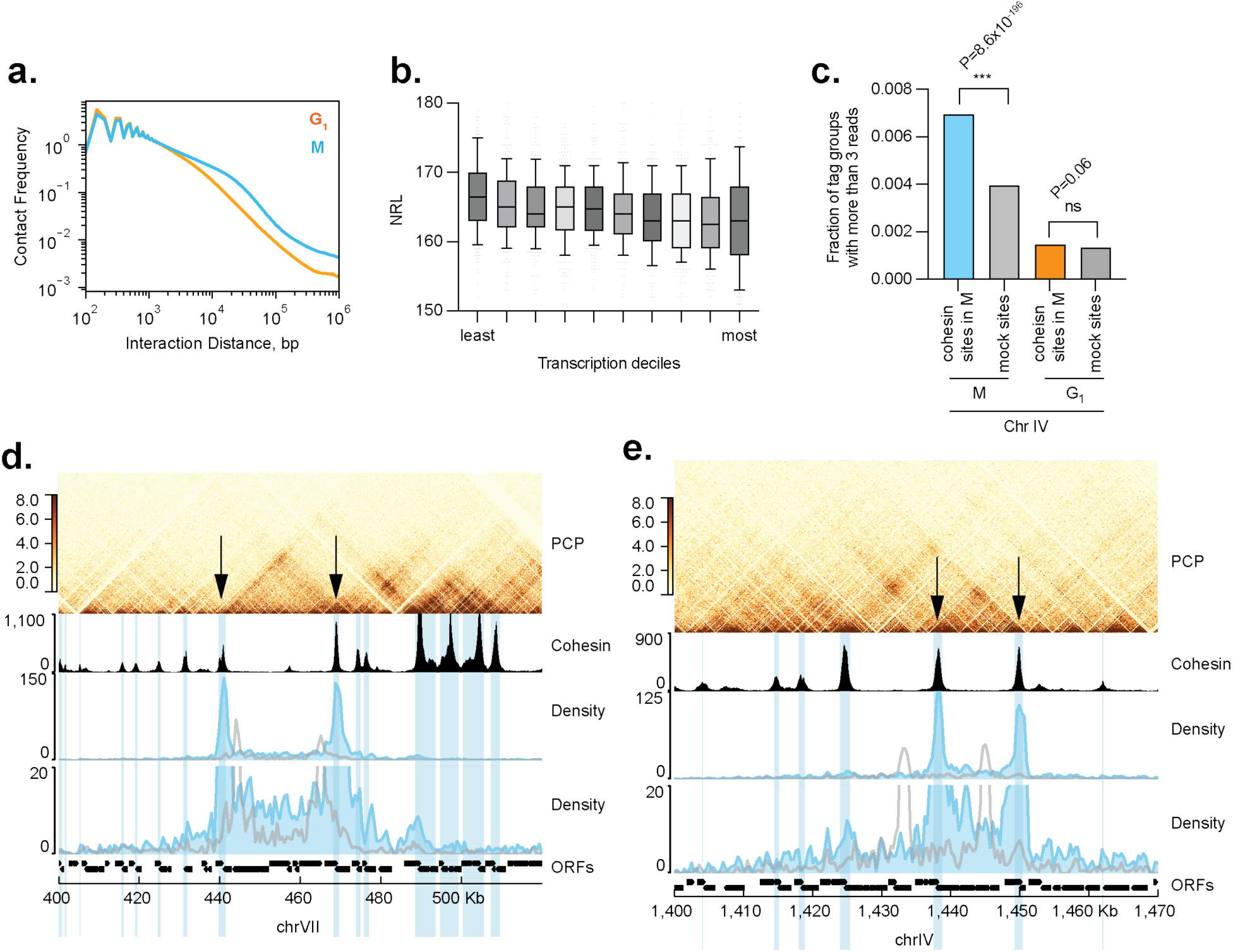
a. Distance decay profiles in G1 and M phase from PCP data. There is an enrichment in long range interaction in M phase as previously described (27, 40). b. Nucleosome repeat length (NRL) in relation to transcription deciles over genes larger than 2Kb. NRL was measured using DANPOS3 on mono-nucleosome signal. The least expressed genes have a higher NRL. Median, Box and Whiskers 10-90 percentiles. c. Fraction of tag groups having reads in at least 3 cohesin binding sites vs mock sites in mitotic chromatin on chromosome IV. Comparison is also made for multi-way contacts at the same sites in G1 arrested cells. Fisher exact test is used to calculate p-value. d. Cohesin enrichment sites participate in multi-way long-range interactions. Selecting for tag groups that contain reads in 2 cohesin binding sites (black arrows) reveals complex multi-way interactions. Cohesin ChIP signal is in black (40). Tag groups that contain reads in 2 adjacent cohesin binding sites (in blue) compared to tag groups that contain reads in 2 adjacent mock sites of same size (in grey). Lower panel is a zoom of the upper intermediate panel. e. Same as in d. for another locus.

**Fig. S7.**
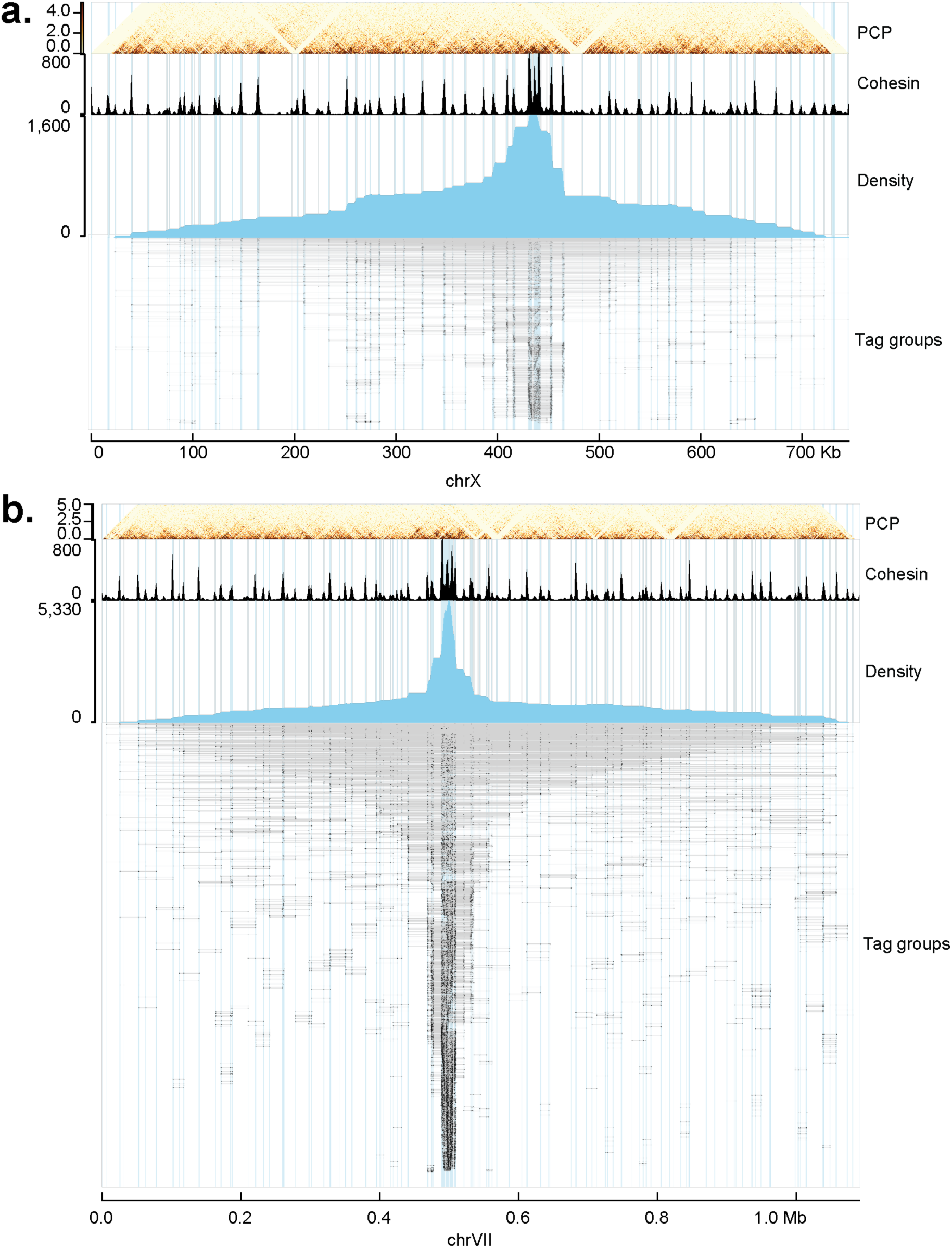
a Estimation of relative chromatin compaction levels across chromosomes. Tag groups including at least one read in 3 different cohesin binding sites are mapped along chromosome IV. Cohesin ChIP signal is shown in black (40). Blue area shows the density profile of the reads represented on the lower panel in which each line corresponds to a tag group, thick black lines are mapped reads, connectivity of each tag group is shown as grey lines. b Same as in a. for chromosome VII.

**Fig. S8.**
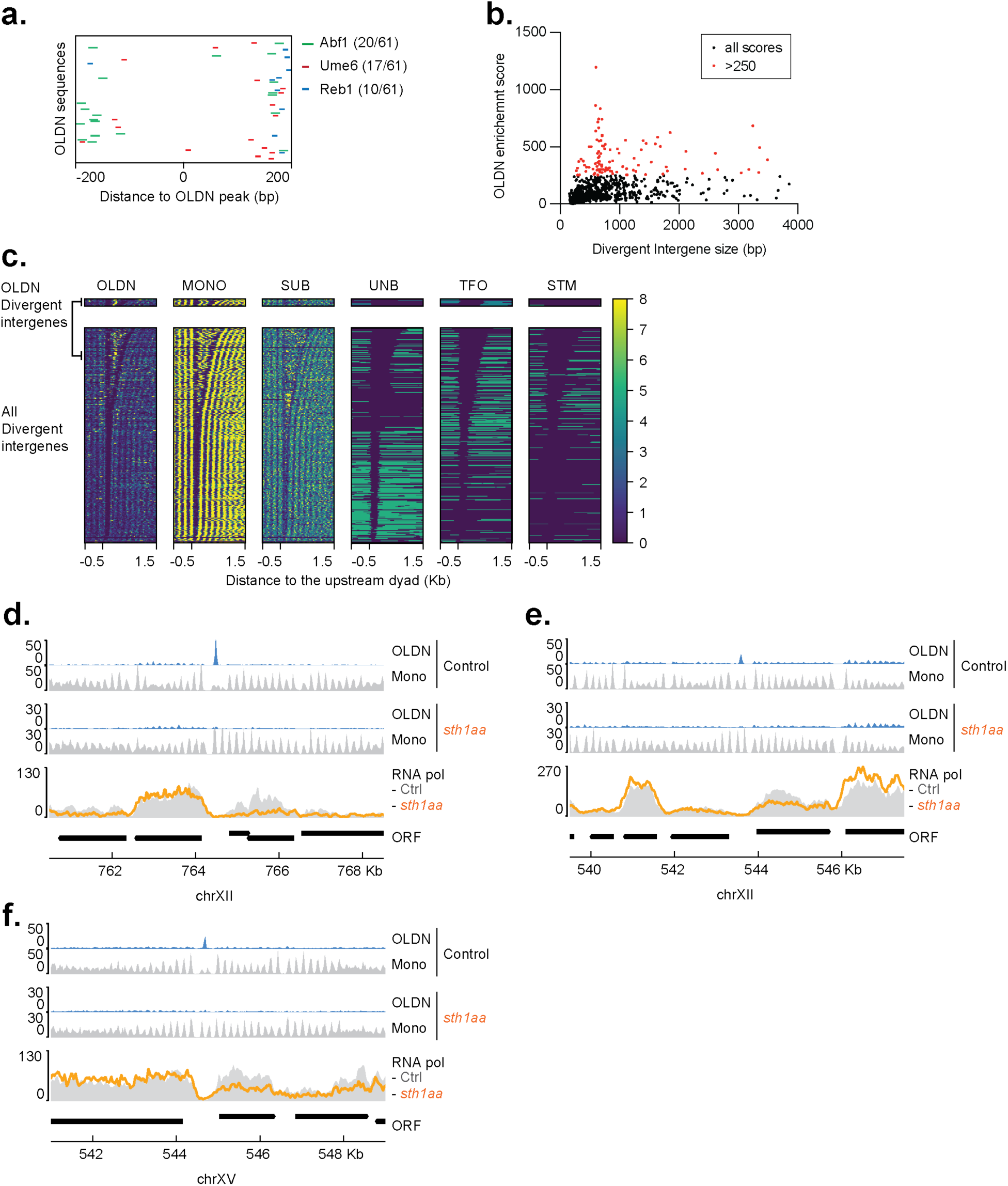
a. Position of transcription factor binding motifs Abf1 (green), Ume6 (red) and Reb1 (blue) centered on the midpoint of the OLDN for 61 sites. b. OLDN enrichment score versus divergent intergene size indicates that OLDN are often found at divergent promoter pairs of distinct sizes. c. Metaplot of size selected PCP data mapping to divergent gene promoter pairs. All divergent intergenes sorted by size: OLDN (210 bp - 270 bp), SUB (20bp - 110bp), (Mono 130-190bp). OLDN are enriched in a certain divergent intergene size. Promoter classes are described in (39). d. (e) (f) Comparison of footprint positions and size distribution in control (top panel) and RSC depletion (*sth1aa*, lower panel) at representative locus. Nucleosome tracks of OLDN size range are blue (210 bp < insert size < 270 bp) and mono-nucleosomes in grey. RNA polII ChIP in both conditions (38) is shown in the bottom panel.

